# Complement C3d enables protective immunity capable of distinguishing spontaneously transformed from non-transformed cells

**DOI:** 10.1101/2024.07.31.606044

**Authors:** Jeffrey L. Platt, Chong Zhao, Jeffrey Chicca, Matthew J. Pianko, Joshua Han, Stephanie The, Arvind Rao, Evan Keller, Mayara Garcia de Mattos Barbosa, Lwar Naing, Tracy Pasieka-Axenov, Lev Axenov, Simon Schaefer, Evan Farkash, Marilia Cascalho

**Affiliations:** Department of Surgery, University of Michigan, Ann Arbor, MI, 48109, USA; Department of Microbiology and Immunology; University of Michigan; Ann Arbor, MI, 48109, USA; Division of Hematology/Oncology, Department of Internal Medicine, University of Michigan, Ann Arbor, MI 48109, USA; Computational Medicine and Bioinformatics, University of Michigan, Ann Arbor, MI, 48109, USA, University of Michigan, Ann Arbor, MI, 48109, USA; Cancer Data Science Shared Research Core of the University of Michigan Rogel Cancer Center; Department of Urology and the Biointerfaces Institute, University of Michigan, Ann Arbor, MI, 48109, USA, University of Michigan, Ann Arbor, MI, 48109, USA; Department of Pathology, University of Michigan, Ann Arbor, MI, 48109, USA; Graduate Program in Molecular and Cellular Biology; Department of Biology, Division of Genetics, Nikolaus-Fiebiger-Center for Molecular Medicine, Friedrich-Alexander-Universität Erlangen-Nürnberg (FAU), Erlangen, Germany

**Author notes:** Corresponding authors. Marilia Cascalho, Address: Medical Science Research Building I, A520B, 1150 West Medical Center Drive, Ann Arbor, Michigan 48109-5656; Jeffrey L. Platt, Address: Medical Science Research Building I, A510E, 1150 West Medical Center Drive, Ann Arbor, Michigan 48109-5656.

## Abstract

Immune-surveillance depends in part on the recognition of peptide variants by T cell antigen receptors. Given that both normal B cells and malignant B cells accumulate mutations we chose a murine model of multiple myeloma to test conditions to induce cell-mediated immunity targeting malignant plasma cell (PC) clones but sparing of normal PCs. Revealing a novel function for intracellular C3d, we discovered that C3d engaged T cell responses against malignant plasma cells in the bone marrow of mice that had developed multiple myeloma spontaneously. Our results show that C3d internalized by cells augments immune surveillance by several mechanisms. In one, C3d induces a master transcription regulator, E2f1, to increase the expression of long non-coding (lnc) RNAs, to generate peptides for MHC-I presentation and increase MHC-I expression. In another, C3d increases expression of RNAs encoding ribosomal proteins linked to processing of defective ribosomal products (DRiPs) that arise from non-canonical translation and known to promote immunosurveillance. Cancer cells are uniquely susceptible to increased expression and presentation of mutant peptides given the extent of protein misfolding and accumulation of somatic mutations. Accordingly, although C3d can be internalized by any cell, C3d preferentially targets malignant clones by evoking specific T cell mediated immunity (CMI) and sparing most non-transformed polyclonal B cells and plasma cells with lower mutation loads. Malignant plasma cell deletion was blocked by cyclosporin or by CD8 depletion confirming that endogenous T cells mediated malignant clone clearance. Besides the potential for therapeutic application our results highlight how intracellular C3d modifies cellular metabolism to augment immune surveillance.

**One Sentence Summary:** We show that intracellular soluble fragment 3d of complement (C3d) induces regression of spontaneous multiple myeloma in mice reducing tumor burden by 10 fold, after 8 weeks. C3d enables cell-mediated immunity to target multiple myeloma clones sparing non-transformed polyclonal B cells and plasma cells with lower mutation loads. We show that C3d increases the expression of ribosomal subunits associated with the translation of defective ribosomal products (DRiPs). C3d also decreases expression of protein arginine methyl transferase (PRMT) 5 which in turn relieves E2f1 repression increasing the expression of Lnc RNAs and derived peptides that evoke anti-tumor cellular immunity. The approach increases MHC-I expression by tumor cells and generates a CMI response that overcomes tumor immune-evasion strategies.

**Significance:** Tumors are immunogenic in part because of somatic mutations that originate novel peptides that once presented on MHC engage cell-mediated immunity (CMI). However, in spite of the higher mutation load most tumors evade immunity. We discovered that a component of the complement system (C3d) overcomes tumor immune evasion by augmenting expression of ribosomal proteins and lncRNAs linked to the presentation of novel peptides by tumor cells. C3d induced CMI targets cancer cells sparing non transformed cells uncovering a novel function for complement in immune surveillance.

## INTRODUCTION

Development of malignancy has been known to involve stepwise transformation fueled at each step by more or less random mutations that alone or in combination originate immunogenic peptides. Ironically those same processes are involved in the maturation of B cells responding to an antigen. The immunoglobulins encoding the B cell receptors accumulate mutations that when selected by antigen cause the growth and accumulation of B cells with receptors with variable regions that can differ from their germline counterparts by more than 20% ^1^. Furthermore, peptides derived from mutated Igs are immunogenic ^2^. An important question and the central goal of the research reported is what mechanisms allow tumors but not non-malignant cells to evoke cell-mediated immunity (CMI) segregating protective from auto-immune responses and whether such segregation can be exploited for immunotherapy without causing the depletion of healthy cells.

Complement has been implicated both in the development and in the control of malignancy ^3-7^. Conditions favoring mutation of malignant cells and conditions associated with malignancy such as decreased acidic heparan sulfate proteoglycans dysregulate complement in part owing to alterations in the binding and function of complement regulatory proteins such as factor H ^8-10^. Hypoxia, common in solid tumors also results in increased C3 turnover which eventuates in the generation of C3d making that protein available the vicinity of transformed cells and in draining lymph nodes. C3d covalently bound to antigen has been known to stimulate antibody responses ^11,12^ but unbound C3d was long considered a biologically inactive, non-toxic byproduct and hence its potential for use in generating tumor immunity was not pursued. However, we found, quite by accident that when C3d is expressed in a tumor, the immune system rapidly acquires ability to recognize and respond to the trace amounts of neo-antigen generated by many tumors ^13^. Because prior research tested anti-tumor immunity against adoptively transferred cells we were not able to discern specificity against malignant versus non-malignant clones, only absence of generalized auto-immunity.

We hypothesized that complement products might facilitate segregation of immunity against malignant cells from immunity against healthy cells and tested the hypothesis in multiple myeloma, a tumor of plasma cells, because there are currently no therapies that differentiate between malignant and non-malignant plasma cells. To test whether CMI could selectively recognize and control transformed cells at different stages of malignant transformation we used a mouse model that develops spontaneous multiple myeloma recapitulating the human disease development as well as normal B cell development ^14^. These mice harbor human c-*Myc* that is expressed upon reversion of a STOP codon in B cells undergoing somatic hypermutation, the Vk*MYC mouse ^15,16^. The Vk*MYC mouse spontaneously develops monoclonal gammopathy and multiple myeloma.

To test the impact of C3d on clonal-specific immune surveillance we manufactured a C3d peptide linked in cis to a peptide-internalization domain and administered the recombinant C3d protein to Vk*MYC mice with monoclonal malignancy. The results presented here show that administration of C3d either at the time multiple myeloma might begin or at later time enables protective CMI to develop and clear highly mutated clones sparing non-transformed B cells and plasma cells owing to the activation of long-non coding RNAs and increased expression of ribosomal RNAs linked to expression of defective ribosomal products by malignant cells.

## RESULTS

### C3d decreases paraproteinemia in mice with spontaneous monoclonal gammopathy

To explore whether C3d could induce CMI against malignant plasma cells and/or pre-malignant precursor cells and hence evoke immune surveillance, we tested whether administration of a recombinant C3d protein, could modify the development and early course of monoclonal gammopathies. We reasoned that if malignant plasma cells are immunogenic, C3d might enhance anti-tumor CMI. On the other hand, if multiple myeloma and premalignant conditions are not immunogenic C3d might not succeed.

To test the impact of soluble C3d on clonal-specific immune surveillance we produced a C3d peptide linked to a peptide-internalization domain (Figure S1A) and administered the recombinant C3d protein in mice that develop spontaneous monoclonal malignancy, the Vκ*MYC mice. Vκ*MYC mice express human c-*Myc* under the control of Ig promoter and enhancer in B cells, following B cell antigen activation and reversion of a stop codon by somatic hypermutation ^15^. Tracing the development of pre-malignancy and evolution of multiple myeloma is facilitated by detection of clonotypic M protein, a monoclonal Ig detected in blood and marker of the premalignant monoclonal gammopathy of undetermined significance (MGUS), smoldering multiple myeloma (SMM) ^17^ and ultimately multiple myeloma. Drug efficacy studies in the Vκ*-MYC model predict single-agent drug activity in multiple myeloma, with a positive predictive value of 67% (4 of 6) for clinical activity, and a negative predictive value of 86% (6 of 7) for clinical inactivity ^16,18^.

To accelerate the development of the monoclonal gammopathy Vk*-MYC transgenic mice (between 5 and 8 months of age) were immunized repeatedly (3 times separated by 30 days) with 4-hydroxy-3-nitrophenyl-OVA, NP-OVA, a hapten-carrier conjugate known to induce germinal center reactions, until development gammopathy and ultimately multiple myeloma were evident from quantification of IgG and clonotypic M protein, recapitulating disease progression in humans ^15,16^. We administered 20 μg of C3d protein by intra-tibial injection to Vκ*-MYC mice that had a clear monoclonal Ig band. Untreated mice were injected with the buffer (PBS) without C3d. Because monoclonal gammopathy and multiple myeloma develop at different ages in Vκ*-MYC (as in humans), mice were paired according to age, baseline total and monoclonal IgG concentrations, and one of each pair was either treated with recombinant C3d or injected with PBS, that is left untreated. The time course of treatment, 3 immunizations in the space of 2 months, followed by intratibial C3d injection as soon as an M protein was apparent, and endpoint 2 months later, took in consideration the time that it takes to develop an adaptive immune response and the half-life of IgG to optimize the likelihood of observing a reduction in the M protein concentration following treatment.

We found that a single C3d treatment averted the increase in serum IgG and M protein in mice of the same age and equivalent tumor burden at the beginning of the experiment. In untreated mice, the serum IgG increased 49% during the ensuing two months (Figures 1A and 1B) and the M protein increased in every mouse injected with PBS (Figures 1C and 1D, Figures S1B and S1C). In contrast, after treatment with C3d for 2 months, the concentration of IgG in serum decreased, on average, 32.5% (Figures 1A and 1B) and the M protein bands detected by serum protein electrophoresis at the time of injection became indistinguishable from broader Ig bands (Figures 1C and 1D, Figures S1B and S1C). Thus, over two months, treatment with C3d selectively decreased paraproteinemia, while allowing or possibly favoring production of polyclonal IgG. The result is at least consistent with the possibility that C3d promoted development of immunity that selectively impaired Ig production by transformed cells.

**Figure 1:**
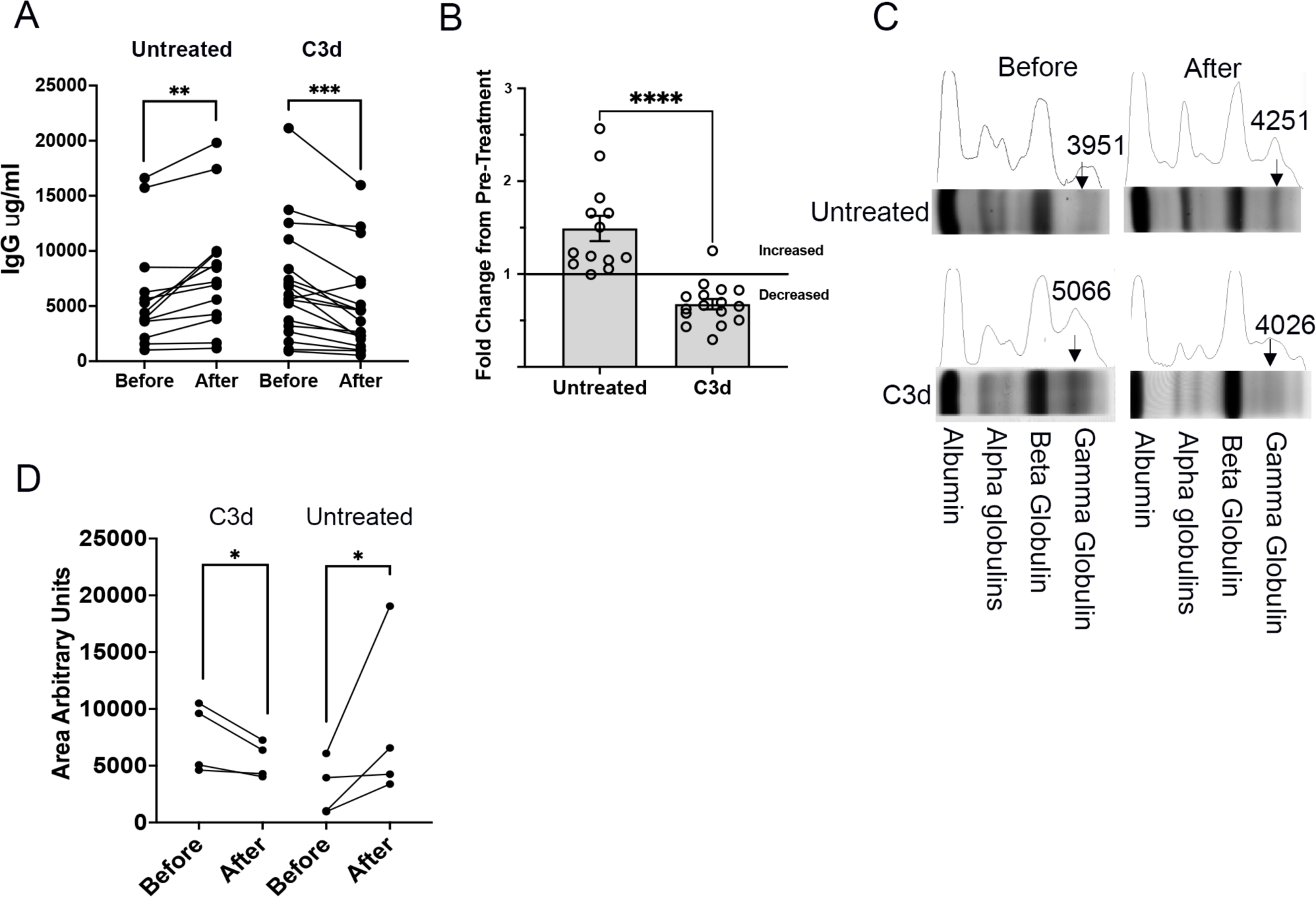
C3d injection of Vk*myc mice decreases paraproteinemia. Vk*myc mice older and younger than one year (N>12), with detected M-protein, were injected with 20 microg C3d peptide intra-tibia two months before analysis. A and B. Shown is the change in the blood IgG concentration in Vk*myc mice following C3d or control injection. C. M protein (arrows) was detected by SPEP before and at the end of treatment. M protein, seen as a single narrow band in the gamma region of the gel decreases in mice treated with C3d. Instead, a broad band of polyclonal Ig appears. In control mice the narrow band increases relative to the albumin band. D. Graph shows the area under the globulin curve comparing analysis of the blood proteins before and after treatment with C3d or with PBS. The units are arbitrary and were determined by selecting the respective areas in image J using the average between lower peaks as the bottom line. A. Comparisons were by double tailed paired t test. **, P<0.01;***, P<0.001. B. Comparison was by two tailed Mann-Whitney test. ****, P<0.0001. D. comparisons were by Ratio Paired T test, P=0.03.

### C3d treatment decreases paraproteinemia

To determine whether the decrease in paraproteinemia following treatment with C3d reflected a decrease in tumor burden, we assayed the frequency of plasma cells in bone marrow retrieved from Vκ*-MYC mice 60 days after treatment with C3d or from paired Vκ*-MYC mice that were received PBS only. Vκ*-MYC mice treated with C3d 60 days earlier, had in bone marrow, less than two-fold the number of plasma cells detected by Cytometry by Time of Flight (CYTOF) than PBS injected partners (Figures 2A-2C). Although bone marrow samples exhibit notable variability in cellularity, all treated mice save one had appreciably lower frequencies of plasma cells than untreated partners (Figure 2C). This extent of decrease in plasma cell burden was confirmed by flow cytometry and by ELISPOT of bone marrow derived cells (Figures 2D-2E). The decrease in the frequency of ASCs was not observed in the spleen (Figure 2F) confirming a BM specific decrease. The low frequency of ASCs in the spleen is consistent with a 3 months’ earlier immunization.

**Figure 2:**
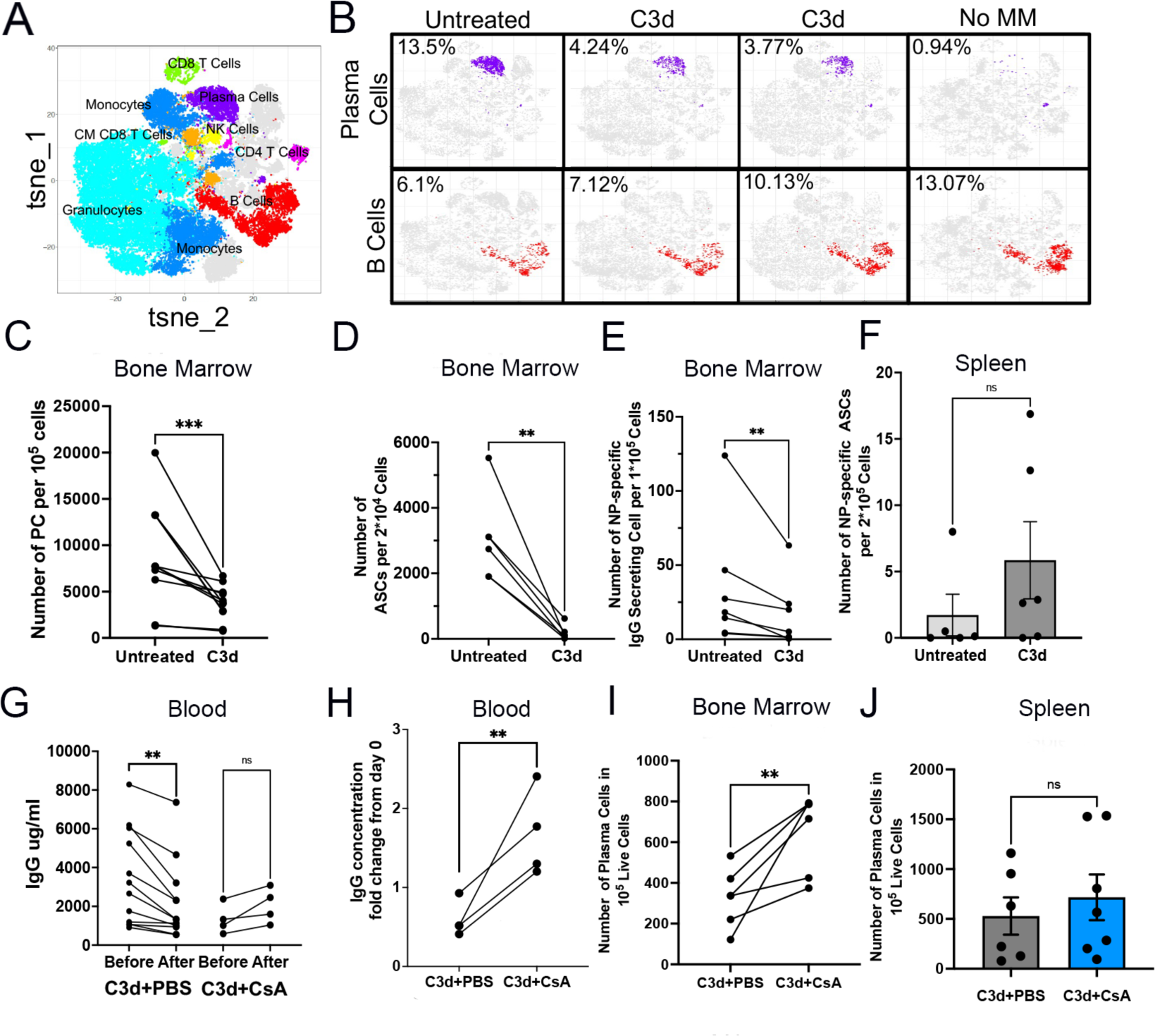
C3d induces CMI that in turn depletes malignant cells. Vk*myc mice older and younger than one year, (N>10), with detected M-protein, were injected with 20 μg C3d peptide intra-tibia two months before analysis. M protein was detected by SPEP. To accelerate the development of Multiple Myeloma, Vk*myc mice were immunized three times with 4-hydroxy-3-nitrophenyl (NP) conjugated to ovalbumin (NP-OVAL) once every month starting 3 months before treatment with C3d. A. Tsne plots of mass cytometry (CYTOF) indicating the various cell clusters. B. Typical CYTOF tsne plots of plasma cell clusters in purple and B cells (not plasma cells) in red. CYTOF analysis of a representative C57BL/6 mouse of comparable age to the plots of the Vk*myc mice is shown in the far right (Control without MM). C. Frequency of plasma cells detected by CYTOF of bone marrow. D-F. ELISPOT analysis of IgG antibody secreting cells (D) or of NP-specific IgG ASC (E) in BM, or of NP-specific B cells in the Spleen, SP (F), 2 months after C3d peptide or with PBS injection intra-tibia. G-J. Vk*myc mice treated as explained above were divided in two groups seven days after C3d injection. One group was treated with Cyclosporin (CsA) administered at a concentration of 3 mg/kg twice weekly for 7 weeks until end point, 2 months after C3d injection. G and H. Serum IgG concentration before and after treatment. I-J. Number of plasma cells determined by FACS counting CD19-, B220lo, CD138+ cells in 100,000 live cells in the bone marrow (I) or in the spleen (J). Analyses in C-E and G-I were by double tailed paired t test. **, P<0.01;***, P<0.001. F and J. Comparison was by two tailed Mann-Whitney test. Significance was considered if P<0.05.

The decrease in frequency of bone marrow plasma cells was observed both in elderly mice (>1 year of age) with frank manifestations of multiple myeloma and in younger mice (0.75 years of age) that had not developed full multiple myeloma (IgG < 5mg/ml) (Figure S2 A-C). These results suggest the impact of C3d treatment on paraproteinemia reflects at least in part a decrease in tumor burden and that the decrease can impact partly transformed pre-malignant as well as malignant plasma cells.

### CMI targets malignant clones and spares non-malignant clones

From theoretical and practical perspectives, it would be important to know whether the decrease in serum IgG and in the frequency of plasma cells, reflected a specific action on multiple myeloma cells and possibly pre-malignant plasma cells or, a broader action that impacted normal plasma cells, normal hematopoietic or non-hematopoietic cells. As mice treated with C3d appeared healthier and better groomed than untreated partners, treatment with C3d generated no obvious systemic toxicity. Moreover, treatment with C3d had no discernable impact on frequencies of hematopoietic cells (Figure S2D), suggesting the impact could be limited to cells of B cell lineage. The more difficult and discerning question however is whether treatment with C3d impacts non-malignant cells of B cell lineage, as normal B cells and especially memory B cells and plasma cells accumulate mutations to the same extent as multiple myeloma cells ^19^ and display cell surface phenotypes similar to phenotypes of malignant plasma cells.

We next asked whether C3d selectively targets transformed plasma cells or both, transformed and non-transformed plasma cells. Since multiple myeloma and pre-malignant gammopathies were induced in Vκ*-MYC mice by immunization of mice with NP-OVA (which would selectively induce hMYC in NP-specific B cells), this question might be approached in part by comparing frequencies of NP-specific ASC in bone marrow where MM cells tend to accumulate, and spleen, where naïve and activated B cells accumulate. Vκ*-MYC mice treated with C3d had, on average, 16-fold fewer NP-specific antibody-secreting cells (ASC) in bone marrow than untreated Vκ*-MYC mice (Figure 2E). However, treatment with C3d did not suppress T cell and B cell responses to NP-OVA, as NP-specific ASC in the spleens of treated mice were as frequent as in in spleens of untreated mice even if their frequencies were low owing immunization 3 months earlier (Figure 2F). These results are consistent with the concept that C3d selectively promotes selective clearance of malignant plasma cells.

The notably specific clearance of malignant plasma cells and the sparing of non-malignant plasma cells by treatment with C3d protein suggests the protein might have recruited protective CMI. However, C3d might instead have exerted some yet undefined direct action on malignant plasma cells or it might have promoted antibody-mediated clearance of plasma cells, consistent with functions of C3d conjugated antigen ^11,12^. Our results showing that treatment of Vκ*-MYC mice with cyclosporine, twice weekly beginning 7 days after C3d injection, inhibited C3d-induced decrease in tumor load (Figures 2G-I) while having no effect in splenic plasma cells (Figure 2J), are consistent with the concept that C3d protein recruited cell-mediated immunity to surveil and clear malignant clones. Depletion of CD8 T cells in a limited group supports the concept that C3d mediated clearance of plasma cells depends on cell-mediated immunity (Figures S1B and S1C).

To probe C3d-induced CMI on normal and malignant plasma cells we used single cell RNA sequencing (scRNA seq) analysis to compare bone marrow cells from Vκ*-MYC mice treated with C3d with bone marrow cells from Vκ*-MYC mice that had not received C3d, using 10X genomics technologies. For this analysis, plasma cells were defined as cells with rearranged *IgH* that expressed *hmyc, sd1c, pdrm1*, or *xbp1* but not *cd19* (Figure S3A-B). After filtering, we obtained a total of 3995 cells from 5 Vκ*-MYC mice treated with C3d and from 5 Vκ*-MYC mice that were not treated with C3d. Based on criteria introduced by Croucher et al. ^20^ malignant and non-malignant cells are distinguishable according to the expression of a set of genes associated with malignancy or with non-malignancy. We compiled scores reflecting malignant genes or non-malignant genes expression (see methods). Malignant cells were defined as those with malignant gene expression score in the top 20% and non-malignant gene expression score in the lower 20%. In contrast, non-malignant cells were defined as those with malignant gene expression score in the bottom 20% and non-malignant gene expression score in the top 20%^20^ (Figures 3A-C, Tables ST1-ST5). Only 3% of the plasma cells retrieved from bone marrow from C3d treated Vκ*-MYC animals were malignant, while 28.3% of the of the plasma cells retrieved from bone marrow of mice not treated with C3d were malignant (Figure 3D). Conversely, benign plasma cells were 19-fold more frequent than malignant clones in C3d-treated animals while malignant and non-malignant plasma cells were detected in approximately equal frequencies in Vκ*-MYC mice that were not treated with C3d (Figures 3D). Mice that received PBS had more larger clones than mice given C3d (Figure 3E). Moreover, one in 5 Vκ*-MYC mice treated with C3d had no detectable malignant cells among 651 single cells analyzed, and only 0.7% plasma cells in the bone marrow.

**Figure 3:**
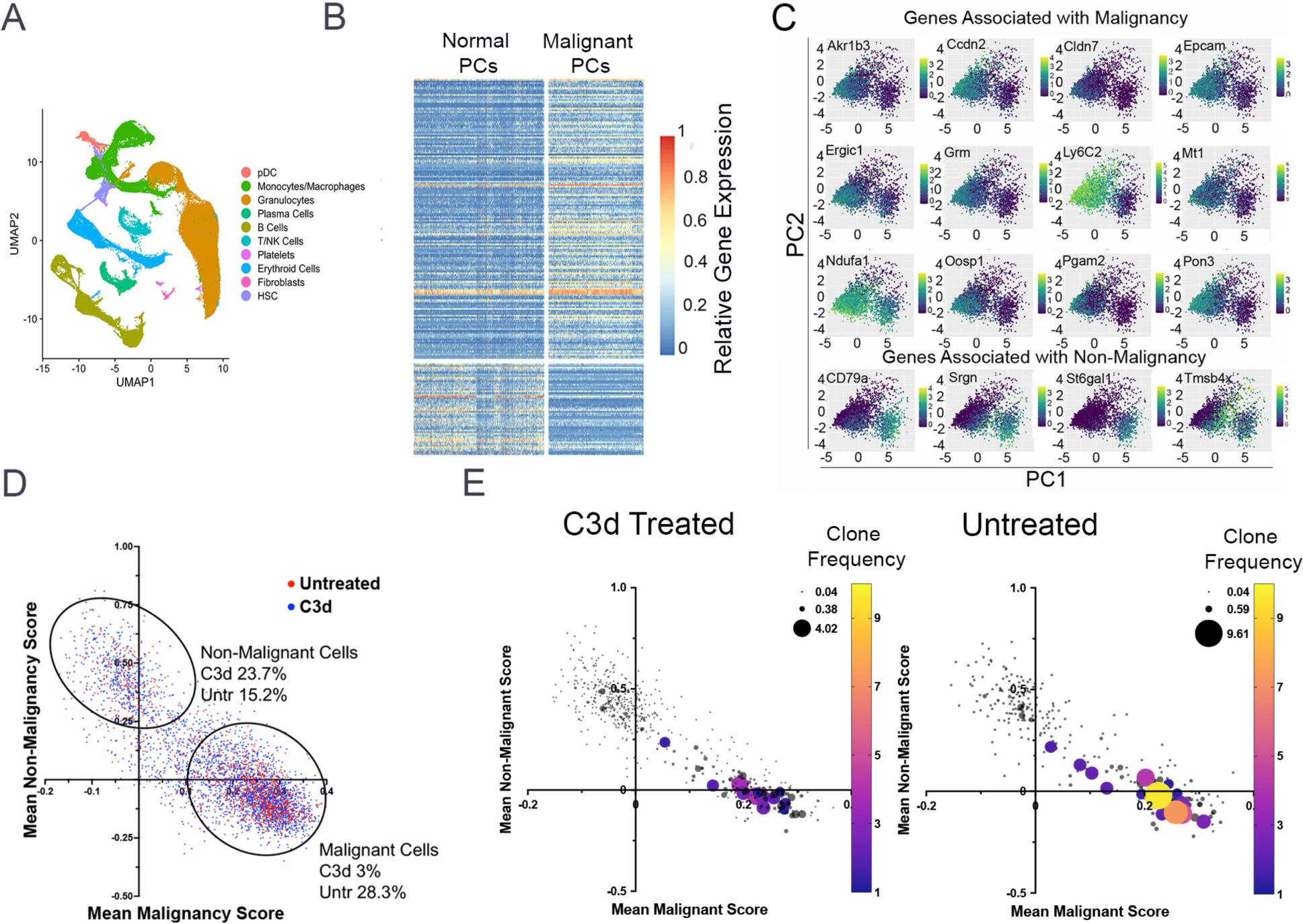
C3d decreases malignant plasma cells sparing normal plasma cells. Vk*myc mice (N=11) were injected with 20 microg C3d peptide or with PBS intra-tibia two months before analysis. Malignancy and non-malignancy associated gene expression was analyzed in 3995 individual plasma cells obtained from the bone marrow. A. UMAP plots of sc RNA gene expression depicting the different cell populations analyzed. B. Graph depicts the differential expression of a set of malignancy associated or non-malignancy associated genes in the upper and lower percentil 20 for each category and in each group of mice. Each horizontal line represents a gene, each vertical line represents a cell. C. Graphs show PCA analysis of the expression of selected genes in the upper and lower percentil 20 associated with malignancy or non-malignancy in all the plasma cells analyzed from both C3d treated and untreated mice. The genes chosen contribute most of the difference between malignant and non-malignant cells. D and E. Clone relative expression of malignant or non-malignant genes in the upper and lower percentil 20, aggregated into scores. Two dimensions plots depicting gene expression scores from treated and untreated mice. (D). The size and color of the circles represent the number of copies of each clonotype defined according to the CDR3 sequence, VH and JH identity. (E).

To determine if highly mutated cells were preferentially targeted following C3d administration we studied the bone marrow Ig B cell repertoires of treated and non-treated mice using 10X genomics. Malignant cells are the progeny of activated B cells that accumulate mutations and express Ig class switched isotypes while the majority of B cells in the bone marrow are immature expressing predominantly non-mutated IgM/D. We analyzed the V(D)J H+L sequences obtained by paired NGS of single cells obtained from bone marrow isolates from five Vκ*-MYC mice treated or from five Vκ*-MYC mice not treated with C3d, 8 weeks after treatment. Analysis shows that C3d treatment decreased the frequency of Ig class-switched B cells in the bone marrow and increased the frequency of IgM expressing cells by 2 fold consistent with specific targeting of malignant plasma cells that predominantly express IgG (Figure 4A). We did not observe significant differences in the VH repertoires of bone marrow B cells in mice treated or not with C3d suggesting that C3d spares normal B cell development that is the major source of repertoire diversity in the bone marrow (Figures 4B-4D and S4). We also observed that IgH CDR3 length distribution obtained from mice that received PBS was skewed likely owing to over representation of certain malignant clones (Figure 4E). Furthermore, analysis of V-J recombination of all IgH chains by Circos plots showed that there are fewer enlarged clones that utilize the same VH and JH gene segments in mice treated with C3d relative to untreated mice (Figures 4C-D). C3d treatment decreased the frequency of Ig clonotypes expressing VH mutated above 2% by almost 6 fold (Figure 4F) and decreased the frequency of VH mutations for cells expressing IgA and IgG1 isotypes (Figure 4G). These findings are consistent with C3d-dependent CMI, selective elimination of malignant clones that have highly mutated Ig, express Ig class switched isotypes and are found in multiple cells while preserving normal B cell development of cells expressing mostly unmutated IgM and a diverse V(D)J repertoire.

**Figure 4:**
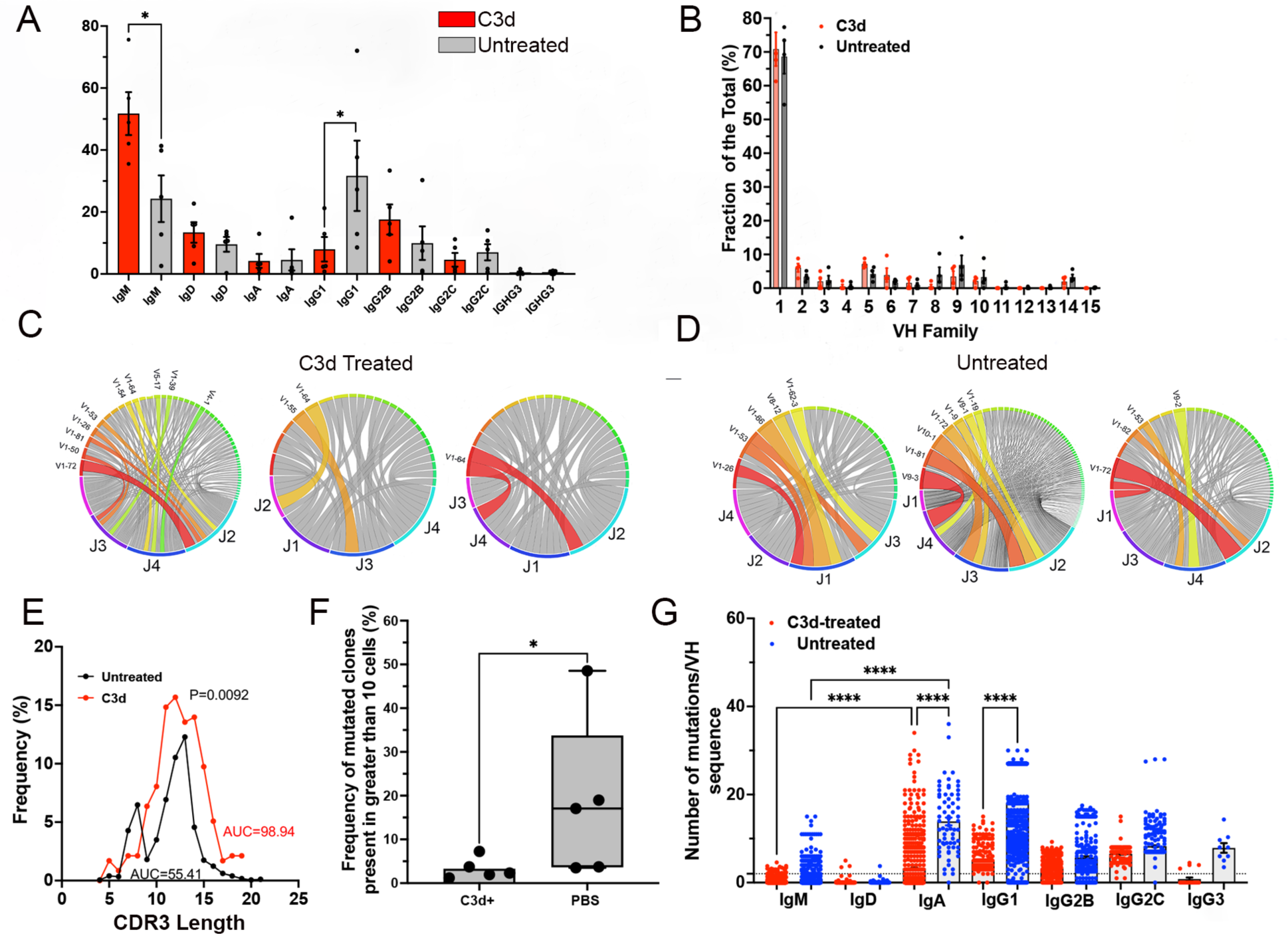
C3d treatment decreases the frequency of highly mutated malignant B cell clones and Ig class switched B cells in the BM but increases the frequency of normal maturing B cells. Vk*myc mice were injected with 20 microg C3d peptide or with PBS intra-tibia two months before analysis (N=8). Single cell V(D)J sequences were obtained using the 10X genomics kits followed by NGS. IgH+L sequences were further analyzed using the ARGalaxy immune repertoire pipeline as well as the NCBI Ig Blast and IMGT suites. A. Frequency distribution of Ig isotypes in 5 Vk-MYC* mice treated or not (Unt) with C3d. Comparison of the frequencies of IgM producing clones in the BM by Mann Whitney test of mice treated or not with C3d indicated P= 0.0157. IgG1 frequencies also differed according to the treatment (P=0.0476 Mann Whitney test). B. IgVH family frequencies in B cells in the bone marrow of Vk-MYC* mice treated or not (Unt) with C3d. Differences in the frequencies according to treatment did not reach significancy by multiple Mann Whitney tests. C and D. Typical Circos analysis (ARGalaxy immune repertoire pipeline) of the V-J recombination in H chains obtained from bone marrow of mice C3d treated (C) or not (D) with C3d. The ribbons link each VH to their respective JH and the ribbon thickness is proportional to the number of sequences that express that combination. Around the circle the colored bars reflect each JH or VH. Figure shows there are fewer colored ribbons in C3d treated mice (C) as compared to untreated (D) mice that have wider and/or more frequent VH-JH ribbons, indicative of clonal skewing. E. CDR3 AA length distribution curves obtained from bone marrow B cells of mice C3d treated or not with C3d. Untreated CDR3 lengths are skewed and show a peak of 8 AA length. Comparison of the curves by the Wilcoxon matched ranks test indicates an exact P value of 0.0092. The area under the curves (AUC) is also indicated. F. Graph shows the frequency of Ig clones (larger than 10 cells) with VH mutation frequencies above 2%. Each dot represents the average frequency of mutated clonotypes in all the V(D)J sequences obtained from one mouse marrow. Data analysis by Mann Whitney test indicated a P= 0.0317. G. Number of mutations per VH region in heavy chain sequences obtained from B cells isolated from the bone marrow of Vk-MYC* mice treated or not (Unt) with C3d. Each dot represents mutation frequencies in VH sequences in one cell. Each clonotype is counted only once. Frequencies were analyzed by ANOVA followed by Kruskal-Wallis multiple comparisons test. Comparisons that yielded P<0.0001 are noted by ****.

To directly test if C3d directed CMI to specific plasma cell clones we tested specific NP-responses in the bone marrow reasoning that since multiple myeloma arises in response to NP-OVA, clonal-specific deletion should result in the decrease of cells expressing the canonical VH1-72 and/or the W33L CDR1 mutation that increases affinity for NP by 10 fold ^21,22^. We found that C3d treated mice had 10.4 fold less VH1-72 expressing cells than mice injected with PBS (Figure 5A). Furthermore, in mice injected with C3d, only 10% of the VH1-72 sequences had the canonical W33L mutation while 56% of the VH1-72 sequences in mice injected with PBS had the W33L mutation (P<0.0001, Fisher’s exact test) suggesting that C3d promotes clone specific CMI (Figures 5B-D). Results also show that clone-directed immunity targeted the largest and most mutated clones (Figures 5E and 5F). The data are consistent with clonal-specific CMI.

**Figure 5:**
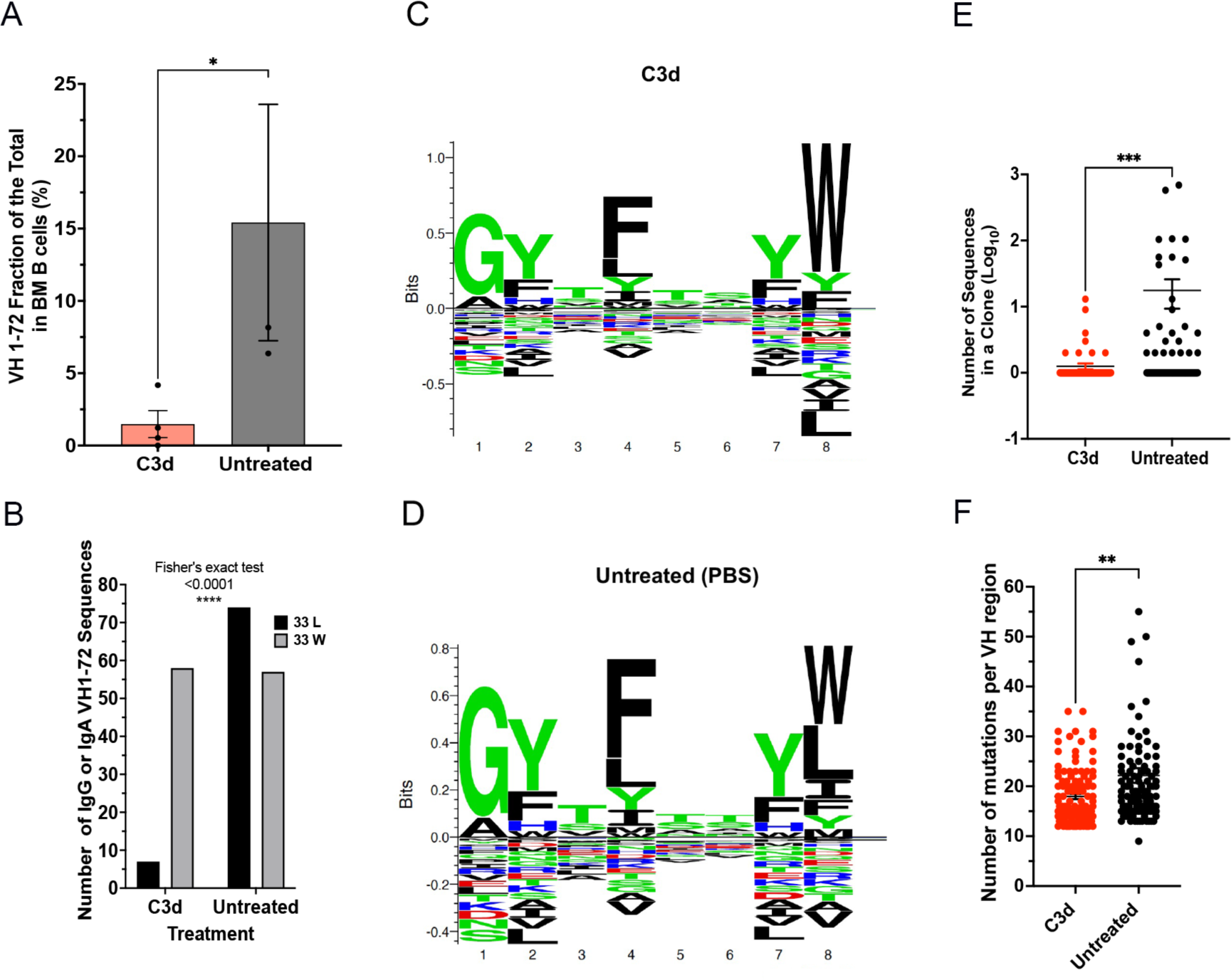
C3d induces clonal-specific deletion of antigen-specific MM cells. Vk*myc mice (N=8) were immunized 3 times with 4-hydrox-3-nitrophenyl(acetyl) (NP) coupled to Ovalbumin with 2 months intervals, starting at 8-10 weeks. Mice were injected with 20 microg C3d peptide or with PBS intra-tibia two months before analysis. Analysis of BM cells was by 10X genomics. A. Graph depicts the frequency of VH1-72 usage in each mouse. VH1-72 is the main VH region recruited by NP antigens. Comparison was by the Mann Whitney test, P=0.0286. B. The number of IgG or IgA VH1-72 sequences in clones obtained from mice treated or not with C3d was compared by Fisher’s Exact test. P<0.0001. C and D. Seq2Logo depicting the CDR1 aminoacid motifs in B cell clones with VH1-72 from mice treated (C) or not (D) with C3d. Stacks represent AA with the size of the letters proportional to their prevalence. Shown is the Kullback-Leibler Logo. AA below “0” depict depleted AA. Apparent is the depletion of L at position 8 (corresponding to AA33 in the VH sequence) in mice treated with C3d. The W33L mutation causes a 10X increase in affinity to the NP hapten. Seq2Logo is available at http://www.cbs.dtu.dk/biotools/Seq2Logo. E. Graph depicts the size of each clone shown as the Log_10_ of the number of sequences in each clone (dot), (C3d N=115; PBS N=110). Clones were defined by 80% similarity in CDR3 AA, same VH and JH. Results were compared by the Mann Whitney test, P=0.0006. F. Number of mutations per VH1-72 regions in heavy chain sequences obtained from B cells isolated from the bone marrow of Vk-MYC* mice treated or not with C3d. Each dot represents the VH mutation frequency/cell. Each clonotype is counted only once. Only VH regions with mutation frequencies above 10% are shown. Results were analyzed by ANOVA followed by Kruskal-Wallis multiple comparisons test.

### C3d increases expression of genes encoding ribosomal proteins and genes encoding long non-coding RNAs that increase MHC-I expression of neo-peptides

C3d treatment caused a profound change in gene expression. scRNA analysis of bone marrow cells revealed that C3d changed the expression of 346 genes in bone marrow cells studied 2 months after treatment, with an adjusted P<0.001 (ST6.1 and ST6.2). To determine how C3d impacted malignant plasma cells we wondered if internalization of C3d by tumor cells altered plasma cell gene expression promoting immunity. scRNA analysis of bone marrow hMyc-positive plasma cells revealed that C3d changed the expression, with adjusted P<0.001, of 58 genes and C3d changed the expression of 70 genes with an adjusted P<0.05 (Figures 6 A-C and ST6.2). Changes in the ribosomal composition determine the extent to which translation of defective ribosomal products “DRiPs” occur ^23^. Because presentation by MHC I of peptides originated from DRiPs are important in immune surveillance ^24^ we asked whether C3d changed ribosomal gene expression to facilitate translation of DRiPs. Results show that C3d increased the expression of 23 of the 80 ribosomal proteins by plasma cells. C3d increased the expression of Rpl genes that encode proteins that form the S60 ribosomal subunit (Rpl10, Rpl22, Rpl23a, Rpl24, Rpl30, Rpl35, Rpl35a, Rpl36a, Rpl37, Rpl37a, Rpl38, Rpl39 and Rpl41) with adjusted P< 0.001. C3d also increased the expression of Rps genes that encode proteins that form the 40S ribosomal subunit (Rps2, Rps3, Rps3a1, Rps4x, Rps15a, Rps21, Rps16, Rps26, Rps27a and Rps29) with adjusted P< 0.001 (ST6.2). Consistent with the increased expression of ribosomal genes gene ontology (GO) term analysis identified “translation and secretion” as the main pathways enhanced by C3d treatment on Plasma cells (Figure 6D). Of the 23 genes encoding ribosomal proteins increased in expression, in plasma cells, by C3d *in vivo*, 13 were also found to be increased following C3d treatment of 18.81 cells in culture, with adjusted P<0.001, (Figure S5B, ST7), compared to cultures that received media alone. C3d is internalized by 18.81 cells suggesting that changes in gene expression are a function of intracellular C3d (Figure S5A). These results are consistent with C3d dependent changes in ribosomal function. Of particular interest, Rps28 expression has been linked to immune-surveillance ^23^ suggesting that intracellular C3d might increase antigen presentation and non-canonical gene translation originating DRiPs for MHC-I presentation.

**Figure 6:**
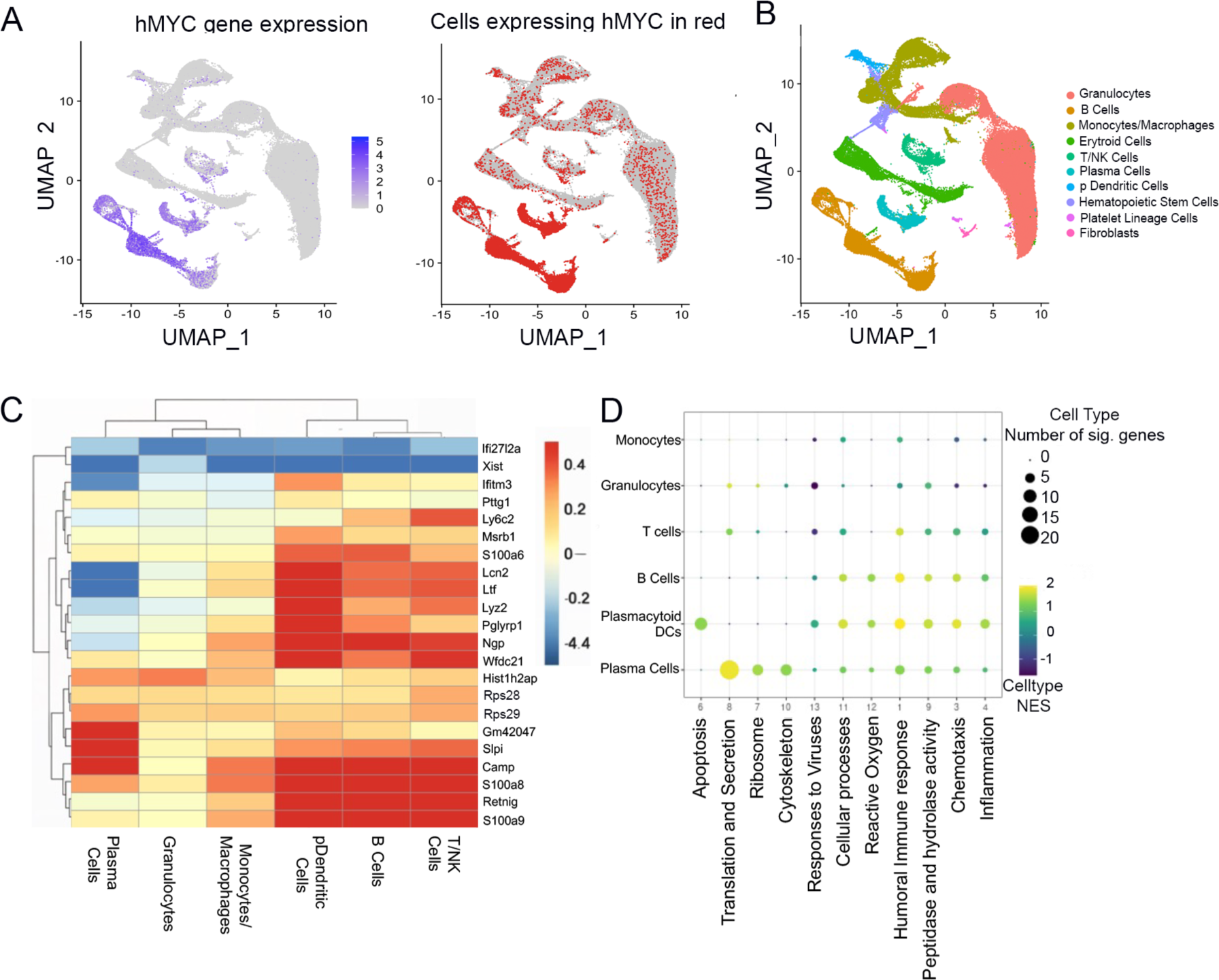
C3d increases expression of genes encoding ribosomal proteins and Long Non-Coding RNAs. Vk*myc mice (N=11) were injected with 20 microg C3d peptide (N=6) or with PBS (N=5) intra-tibia two months before single cell RNA analysis of bone marrow cells. (A and B) UMAP plots of sc RNA gene expression depicting the different cell populations analyzed indicating hMYC expression to identify cells of the B cell lineage that activated hMYC. (C) Genes differentially expressed in C3d treated relative to untreated mice in all the bone marrow cell populations indicated. Scale on the right refers to the Log_2_ fold changes. (D) GSEA pathway analysis of differentially expressed genes in various bone marrow cell populations. Major pathways are identified below and the cell types on the left. The normalized enrichment score (NES) is shown in a colored scale and the number of genes differentially expressed in each pathway are represented in proportion to the size of the circle.

Treatment by C3d increased the expression of long non-coding RNAs shown to provide MHC-I binding peptides^25^. Thus, C3d increased the expression of Gm42047 (adjP=3.19×10^-40^) by plasma cells and to a lesser extent by B cells and plasmacytoid dendritic cells in the bone marrow (Figures 6, S5, and table ST10). Gm42047 expression was also increased after incubation of 18.81 with C3d (Figures 6C and S5B-D, Tables ST6.2 and ST8). Incubation of 18.81 cells with C3d changed the expression of several LncRNAs found by Barczak et al. ^25^ to contribute peptides that bind MHC-I with high affinity. Those included 4930426D05Rik (adjP=4.32*10^-4^), Gm42047 (adjP=1.43*10^-5^), Gm43549 (adjP=3.25*10^-6^) and Gm37499 (adjP=0.006) (Figures S5B-D, S6B, Table ST8).

Many LncRNAs are targets of E2f1, the target of retinoblastoma protein. E2f1 is repressed by methylation by the protein arginine methyl transferase (PRMT)5. Given that C3d promoted changes in the expression of lncRNAs, we asked if the E2f1 pathway was activated by C3d. Analysis of gene expression in 18.81 B cells treated with C3d revealed that along with E2f1, the genes encoding known E2f1 partners were significantly increased in expression following C3d exposure while the gene encoding Prmt5 was decreased (Figure S6). We confirmed that C3d increased E2f1dependent gene expression by quantitative PCR (Figure S5C). The results suggest that C3d activates E2f1 mediated gene transcription of lncRNAs which in turn provide peptides to load on MHC-I engaging T cell-dependent immune surveillance as shown in ^25^. Testing this possibility by incubating 18.81 cells with C3d revealed that C3d increased MHC-I expression by at least 30% starting at 24 hours post-incubation (Figure S7).

Consistent with an activation T cell-dependent responses we found that T cells in bone marrow of Vκ*-MYC mice treated with C3d had increased expression of genes encoding granzyme K and C-C motif chemokine ligand 4 (CCL4). Gzmk encodes granzyme k and the message is expressed by CD8+ T cells in the inflammatory milieu ^26^ (Tables ST9-10). We also found a significant increase in the expression of C-X-C motif chemokine ligand 2, Cxcl2, and Interferon Alpha-Inducible Protein 27-Like Protein, Ifi2712a, in granulocytes in bone marrow of Vκ*-MYC mice treated with C3d compared to granulocytes obtained from controls (Tables ST11-12). CXCL2 is produced by neutrophils and is a potent chemoattractant for these cells. Ifi2712a is involved in apoptosis and its function in granulocytes is not well understood.

## DISCUSSION

We show that C3d enhances CMI that selectively targets transformed cells and selectively spares non-transformed cells revealing a path skewing immunity away from healthy cells. Although a precise understanding of the specificity of protective cell-mediated immunity promoted by C3d polypeptide remains to be determined, the high frequency, rapid development (in 2 months), clear specificity of responses for malignant highly mutated plasma cells and not non-malignant plasma cells, indicate the responses might target the products of multiple mutations of somatic non-Ig genes such as those encoded by long non-coding RNAs and/or an Ig mutation burden clearly increased in malignant plasma cells relative to non-malignant cells. The high frequency of success suggests that immunity either targets antigens for which there are frequent T cell precursors across different mice and or tumor antigens that become immunogenic owing to their prevalence and or context.

The first possibility is supported by findings of C3d-induced ribosomal gene expression associated with increased translation of DRiPs ^23^ and with an increase in expression of long non-coding RNA species found to be associated with production of MHC-I peptide ligands associated with immune surveillance ^25^. The second scenario is supported by a selective decrease in the frequency of VH1-72 plasma cells and in VH1-72 sequences bearing the W33L mutation, consistent with clonal-specific deletion. Although we do not have data addressing the non-Ig mutation burden, the data on Ig mutation burden clearly shows that the frequency of mutated clonotypes in the bone marrow is decreased in mice treated with C3d relative to non-treated mice. Data also show that the size of the clones isolated from bone marrow is decreased, on average, by more than 14 fold, in C3d treated mice relative to non-treated mice, respectively. The C3d mediated decrease of highly mutated clones (with greater than 10 mutations/VH region) and of larger clones suggests that either mutations are the target of immunity owing to the cumulative changes in the Ig sequence or that highly mutated and/or frequent cells are eliminated because of other properties associated with malignancy, e.g., expression of DRiPs or peptides derived from long non-coding RNAs. Current effort is dedicated to test DRiPs directly in cells treated with C3d.

Since both non-malignant and malignant plasma cells accumulate mutations and C3d induces CMI that spares non-transformed cells our results indicate that although mutations drive immunogenicity that does not suffice to drive and/or target CMI. We cannot exclude immunity independent effects on malignant cells such as direct induction of apoptosis or inhibition of malignant progression. Our results indicate that C3d changes gene expression profoundly and although a fraction of the differentially expressed genes occur in various cell types, C3d appears to have most pronounced effects on malignant plasma cells. Thus, C3d increases the expression of genes encoding ribosomal genes and long non-coding RNAs mostly on plasma cells. Because analysis of gene expression in the bone marrow of treated mice occurs at least two months since C3d administration our observations *in vivo* might miss the early changes. To examine the earliest changes in gene expression we performed gene expression analysis in 18.81 B cells cultured in the presence of absence of C3d. Our results indicate that in 48 hours C3d changes the expression of more than 1000 genes with a P<0.001 (Table ST7). Notably, the expression of genes encoding ribosomal proteins and long-non-coding RNAs are among the most increased. The early time point, 48 hours, allowed us to examine which pathways may lead to the enhanced transcription of ribosomal encoding and long non-coding RNA genes. As reported by Barczak et al. ^25^ our results indicate an increase in E2f1 activity accompanied by a decrease in the expression of the gene encoding PRMT5. These findings suggest that early activation of E2f1 precedes engagement of immune surveillance. This result is concordant with prior findings indicating that availability of C3d during priming of an immune response favors the development of immunity against tumor cells owing to properties characteristic of malignancy that include expression of inflammatory mediators as a results of inflammatory cell death, among others ^27^. In contrast, healthy cells may either degrade C3d, or be less likely to produce or present novel peptides, derived from mutated Ig, DRiPs or lnc RNAs, to evoke immunity.

Our results are consistent with findings showing that transformed cells may promote complement activation ^4^ and in turn generate C3d which may either act in the transformed cell, or in the draining lymph nodes during priming. C3d expressing tumor cells block T regulatory cell development in the draining lymph nodes and in this way facilitate tumor-specific CMI ^13^. Beyond the scope of the research related here and yet unexplored is the possibility that soluble C3d interacts with APCs at the time in antigen presenting cells and/or with the developing antigen-specific T cells themselves.

The limited ability of immunity to treat multiple myeloma might be surprising since precursor cells accumulate numerous mutations in immunoglobulin variable region genes and in other genes ^28^ and the frequency of somatic mutation has been linked to the ability of cancer cells to elicit protective immunity ^29-31^ when pathways of tumor-induced immunosuppression are blocked ^32,33^. However, immunotherapy with checkpoint inhibitors has not been successful in multiple myeloma. Expression of immune-checkpoint mediators such as PD-1 on T cells and PD-L1 on were found in malignant plasma cells but unfortunately anti-PD1/anti-PDL-1 antibodies early clinical trials were not successful ^34^. BCMA targeted CAR-T cells frequently induce dramatic complete responses in 39-67% of patients with relapsed and refractory MM ^14,35^, however these therapies are associated with frequent relapses with no evidence of a plateau in the survival curve ^36,37^. These results suggest multiple myeloma escapes immunity even when the targeted antigen (e.g. BCMA) is necessary for plasma cell function ^38^. Furthermore, toxicity limits implementation of the most aggressive treatments in frail subjects. Because C3d engages endogenous T cells the treatment should be considerably cheaper than CAR-T cells, have reduced or hardly any toxicity and since it does not target a single specific tumor antigen C3d is likely to protect against relapses by tumor cells escaping antigen specific immune destruction. Although a fraction of monoclonal gammopathies progress to multiple myeloma over years (1-10% per year), treatment of these conditions to prevent multiple myeloma is currently discouraged by toxicity. An apparent lack of toxicity could also make C3d polypeptide applicable for prevention of multiple myeloma by treatment of such pre-malignant conditions as monoclonal gammopathy of uncertain significance for which there are few therapies. These are outcomes we will be testing.

## MATERIALS AND METHODS

### Study Design

The study goal was to determine how a recombinant C3d peptide changed the natural evolution of spontaneous Multiple Myeloma in murine model. The Vκ*-MYC mice develop spontaneous MM owing to reversion of a stop codon in the hMYC transgene by somatic hypermutation. In this model, disease progression from premalignancy (monoclonal gammopathy of undetermined significance, MGUS) to full MM can be followed by measuring the concentration of the monoclonal IgG in the blood. MM development was induced by repeated immunization with NP-OVA. Mice paired according to state of disease at the start of the study, were injected with C3d peptide or with the solvent intra-tibia. The end point was set at 8 weeks after injection or earlier in case mice showed signs of morbidity. At the study end point M protein and IgG were measured and compared with values prior to treatment in paired mice. We also enumerated plasma cells in the bone marrow and spleen by FACS, CYTOF and ELISPOT. To determine the extent to which C3d treatment spared normal plasma cells we performed scRNA analysis in bone marrow cells. We also determined systemic impact of the disease in the kidneys by pathologic examination. Finally, mechanism of action was tested by blocking T cell function with Cyclosporin A.

### Mice

C57BL/6 wild type (WT) mice were purchased from The Jackson Laboratories. Vk-MYC* mice were obtained from our collaborator, Dr. Leif Bergsagel at the Mayo Clinic, Scottsdale Arizona^15^. Heterozygous WT/Vk-MYC* mice were maintained by breeding C57BL/6 and WT/Vk-MYC* mice. Mice were genotyped by PCR to select Vk-MYC* positive mice according to^15^.

### Immunization and CD8 depletion

To model development of MM, WT/Vk-MYC* mice were immunized with 100 microg of 4-hydroxy-3-nitrophenyl (NP) conjugated ovalbumin (NP-OVAL) (NP(25)-OVAL(#N-5051-100, Biosearch Technology) in complete Freund’s adjuvant. Mice were re-immunized twice, 30 days apart, by administration of NP-OVAL in incomplete Freunds adjuvant, *i.,p*. Immunizations started 3 months before C3d injection. In a few experiments, mice were injected *i.p* with 0.5 mg of anti-CD8 (clone 53-6.7, Rat IgG2a, κ InVivoMAb) or isotype control (clone 2A3, Rat IgG2a, κ, InVivoMAb) antibodies weekly starting one week before injection of 20 u mg of C3d. Anti-CD8 antibody injections continued until euthanasia at 8 weeks post C3d treatment.

### C3d design and administration

Murine C3d sequence encompassing amino acids 1024-1320 was paired with an N-terminal sequence (RKKRRQRRR, cgcaaaaaacgccgccagcgccgccgc) with Cys (C) to Ser (S) substitution at position 1028 to avoid generating a free sulfhydryl group. The fusion sequence was cloned into a baculovirus expression vector by GIBSON assembly and used to transduce Sf9 insect cells. Polypeptide was isolated using NTA Agarose (Qiagen) and eluted with imidazole in PBS. Purity of samples were determined by electrophoresis in 4-20% gradient SDS-PAGE gels. The monomer was detected in the soluble fractions. Aliquots of 1mg/ml were frozen. Twenty micrograms of recombinant murine C3d with a terminal protein transduction domain in 20 microliters of PBS was injected into the tibia. Some mice treated with C3d were also treated with cyclosporine (SIGMA CAS No:59865-13-3, product number PHR1092) 3 mg/kg twice weekly for 7 weeks.

### Flow cytometry

Single cells were acquired from mouse spleen and bone marrow 60 days after C3d treatment and suspended in cell suspension buffer. Cells were counted on a Neubauer chamber and 1 x 10^6^ cells were incubated in Falcon™ Round-Bottom Polystyrene Test Tubes with FVS780 (BD Life Sciences, San Jose, CA) for 10 min to test cell viability. Cells were incubated for 30 minutes at 37°C with the following antibodies: anti-CD19 (clone #1D3), anti-CD8 (clone#88.25) or anti-CD11b (clone #M1/70) were FITC conjugated; anti-CD19 (clone #1D3), anti-B220, CD45R (clone # RA3-6B2), anti-CD4 (clone #GK1.5) or anti-Ly6G (clone #RB6-8C5) -APC conjugated; anti-IgD (clone #11-26c), anti-CD267,TACI (clone #8F10-3), anti-CD279, PD-1 (clone #J43), anti-CD8 alpha (clone #5H10), anti-Ly6C (clone #RB6-8C5), anti-CD5 (clone #53-7.3) -PE conjugated; anti-CD138 (clone #300506), anti-CD21/35 (clone#7G6), anti-CD117, cKit (clone #2B8) conjugated to Prcp-Cy5.5; anti-LY6A/E, Sca-1 (clone #D7) conjugated to PE-Cy7; and anti-IgM (clone #II/41) conjugated to AF488. Following surface staining, cells were fixed and permeabilized with eBioscience^TM^ Fixation-Permeabilization kit (cat# 00-5521-00) and incubated with FITC conjugated anti-Foxp3 (clone # FJK-16s). All antibodies were purchased from Invitrogen and were diluted 1 to 50. The results were read by FACS Canto II (BD Life Sciences, San Jose, CA) and analyzed by FlowJo^TM^ v10.8 Software II (BD Biosciences, San Jose, CA).

### ELISA

Mice sera were acquired from tail veins before and every 2 weeks since C3d treatment until the endpoint for each mouse. Sera were diluted from 1:50,000 to 1:100,000 and concentrations tested by ELISA done according to published methods. The plates were read at 405nm in a BioTek synergy 2 plate reader.

### Serum Protein Electrophoresis (SPE)

Mice sera was obtained from blood retrieved every 2 weeks following immunizations with NP-OVAL. The SPE was performed on a Helena^TM^ QuickGel Chamber according to manufacturers’ instructions using a QuickGgel split beta.

### CyTOF staining and analysis

CyTOF staining and analysis was done at the CYTOF Core of the University of Michigan. Bone marrow cells were acquired 60 days after initiation of C3d treatment. Cells were suspended in PBS (0.04% BSA), on ice and cell quality was confirmed by measuring cell viability by Trypan Blue staining. Cells were incubated were stained with 50µL of antibody cocktails in Maxpar Cell Staining Buffer, according to the Fluidigm protocol followed by the CYTOF Core. Antibodies were purchased from and conjugated by either the Lederer’s Lab of Brigham and Women’s Hospital or by the Fluidigm Corp., San Francisco, CA.

After staining cells were fixed wit 1.6% formaldehyde solution. Fixed cells were incubated with Cell-ID Intercalator-Ir solution at 4 °C overnight, washed with Maxpar Cell Staining Buffer and resuspended in Maxpar Water the following day. The EQ Four Element Calibration Beads were used to normalize data using a bead-based passport specific to the manufactured bead lot. 50,000 cells were acquired per sample on CyTOF Helios system (Fluidigm Corp., San Francisco, CA) at approximately 200-300 events/s. After acquisition, the instrument software applied a signal correction algorithm based on the calibration beads signal to correct for any temporal variation in detector sensitivity.

CyTOF data analysis: Normalized data was generated at the University of Michigan CYTOF CORE laboratory using standard methodologies. Events were pre-processed by FlowJo^TM^ software for Windows (v 10.8) for live single cells. Gating for intact cells was done by selecting events that express both ^191^Ir and ^193^Ir markers on a biaxial plot using the two DNA channels (^191^Ir and ^193^Ir). Exclusion of beads and cell/bead doublets was done according to expression of ^191^Ir in a DNA channel and excluding ^140^Ce signal, a bead-only isotope. Cells were ^191^Ir-positive and ^140^Ce-negative. Gating for live cells was done by excluding ^195^Pt positive events.

Cytofkit^39^ was used for the data merging and analysis. 10000 Cells in each sample were abstracted to merge and data of 30 to 35 markers was transformed by Customized Asinh Transformation. The t-Distributed Stochastic Neighbor Embedding (t-SNE) algorithm was used for dimensionality reduction and cell subset clustering was generated by Rphenograph^40^. Cell subsets were identified by visualizing the expression of cell-specific surface markers.

### ELISPOT

To enumerate antibody-secreting cells, 96-well Filtration Plate MultiScreen HTS HA Sterile Plates (MilliPore Cat#MSHAS4510) were activated with 30 μl/well of 35% ethanol for 30 seconds. For determining Ig secreting cells, plates were coated with 100 μl of 5μg/ml of unconjugated goat anti-mouse Ig (H+L) (4ug/ml, Southern Biotech, Birmingham, AL, Cat#1010-01, RRID:AB_2794121) in PBS over night at 4°C; for determining NP-specific antibody secreting cells, plates were coated with 100μl of 5 µg/ml NP-24-BSA in sodium carbonate (pH 9.6) overnight at 4°C. Coated plates were blocked with 5% BSA in PBS at room temperature for 2 hours, and cells obtained from spleen and bone marrow were 1:2 serially diluted from a maximum of 2×10^5^ cells per well and incubated at 37° C in 5% CO^2^ atmosphere overnight. Antibody secreting cells were detected with alkaline phosphatase (AP)-conjugated antibodies goat anti-mouse IgM (0.5 µg/mL; SouthernBiotech Cat#1020-04, RRID:AB_2794200) or goat anti-mouse IgG (0.5 µg/mL; SouthernBiotech Cat#1030-04, RRID:AB_2794293) for 1 hour at room temperature. The reaction was visualized by subsequent addition of BCIP/5-bromo-4-chloro-3-indolylphosphate/nitro blue tetrazolium substrate (Sigma-Aldrich Cat#B5655-5TAB). The number of spots was determined using a CTL ImmunoSpot S5 UV Analyzer equipped with ImmunoSpot ImmunoCapture and ImmunoSpot Counting software (Cellular Technology Ltd., Cleveland, OH, RRID:SCR_011082).

### Single-cell-RNA analysis

#### Pre-processing

scRNA was obtained from bone marrow cells isolated from 5 C3d treated and 5 non-treated mice, two months after treatment using 10X genomics procedures. Data were processed and analyzed on on UMICH’s Advanced Research Computing (ARC)’s high-computing cluster, Great Lakes. Reads were aligned and quantified using Cell Ranger using a custom reference to account for the human MYC transgene, which the human MYC gene from GRCh38 was added to the GRCm38 genome reference ^41^. The downstream analysis was done with Seurat in R using the filtered count matrices from Cell Ranger. Cells from each sample were filtered based “nCount”, “nFeature”, and percent mitochondrial genes. Seurat’s integration pipeline was then used to process the scRNA-seq^42^. The cells were integrated by run ID since some samples were sequenced on different days. Reciprocal PCA (RPCA) was used in the “IntegrateData” function. 22 principal components (PCs) were used for clustering and generating the UMAP (Figure S3). Clusters were annotated using canonical cell type markers and cluster-specific differential expression (DE) genes from the “FindAllMarkers” function (Tables ST1-2 and Figure S3). Hematopoietic stem cells (HSCs) expressed Cdk6, Kit, and Adgrg1. Fibroblasts expressed Col1a1, Col1a2, Loxl1, and Lum. Erythroid cells expressed Hba-a1 and Hba-a2. Platelets expressed Pf4, Itga2b, Gp1ba, Tubb1, and Ppbp. T and NK cells were found to be in the same cluster, which these cells expressed a combination of Ptrprc, Cd3d, Cd3e, Cd4, Foxp3, Cd8a, Nkg7, Klrg1, and Gzma. B cells expressed Cd79a, Cd19, and Ms4a1. Plasma cells expressed Sdc1, Xbp1, Ccr10, Prdm1, and Ly6c2 and didn’t express any of the B cell markers. Granulocytes mainly expressed Ptprc, Itgam, S100a8, S100a9, Cd33, Ly6g, Prss34, and Mcpt8. Monocytes/macrophages mainly expressed Ptprc, Itgam, Cd68, Adgre1, Cd14, Mrc1, and Ccr2. Plasmacytoid dendritic cells (pDCs) mainly expressed Ptprc, Cd4, Cd68, Siglech, and Irf8 (Figure S3).

To distinguish malignant versus non-malignant plasma cells, we analyzed gene expression in plasma cells defined as cells expressing one or more of each of the following genes: *Sdc1*, *Xbp1*, or *Prdm1.* We calculated malignant and non-malignant scores based on the expression of genes found to be associated with malignancy in the Vk-MYC* mouse model according to Croucher et al.^20^. Malignant and non-malignant scores were determined in each sample using the “AddModuleScore” function in Seurat^42^. Cells with a malignant score in the top 20% and non-malignant score in the bottom 20% were considered the most malignant; while cells with a malignant score in the bottom 20% and non-malignant score in the top 20% were considered non-malignant (Table ST5). We also ran PCA to distinguish malignant versus non-malignant plasma cells (Tables ST 6-7). The frequency of malignant and benign cells in each sample was calculated considering cell unique identifiers. Heatmaps were created using “pheatmap”. We examined a total of 808 clones.

#### Immunoglobulin sequence analysis

Sequences were obtained from 10X genomics from bone marrow cell suspensions. Data were processed and analyzed on analyzed on UMICH’s Advanced Research Computing (ARC)’s high-computing cluster, Great Lakes. FASTA reads were produced by Cell Ranger software (10X genomics) and sequences were submitted to IgBlast to generate AIRR files or to IMGT ^43-46^ to obtain IMGT archive files. The IMGT files were further analyzed with wither the ARGalaxy suite software ^47^ and/or the statistics program at the IMGT site. From the organized data we compiled isotype and somatic hypermutation data and graphs were produced using prism (v9 for Mac).

### C3d incubation of 18.81 cells

Cell lines, C3d+ 18.81 (passage 7) and C3d- (18.81 passage 8) containing a hygromycin B resistance plasmid ^48^ were cultured at 37°C in RPMI (Thermo Fisher Scientific, Cat: 11875093) supplemented with 10% fetal bovine serum (Thermo Fisher Scientific, Cat: A5670701), 1% penicillin/streptomycin (Thermo Fisher Scientific, Cat: 15140122) and L-glutamine (Thermo Fisher Scientific, A2916801), 0.1% 2-metacaptoethnol (Thermo Fisher Scientific, Cat: 21985023), and 200 µg/ml of Hygromycin B (Thermo Fisher Scientific, Cat: 10687010). Cells were plated at 1×10^5^ cells/100µl in wells of a flat-bottom 96 well plated and incubated for one hour at 37C. Following, the incubation, 20 µl of 1mg/ml C3d or PBS with 10% glycerol was added to wells. This was repeated every 24 hours for two additional days. Total C3d incubation times were 24, 48, and 72 hours.

Cells were harvested and RNA obtained in triplicate per condition using Qiagen RNeasy© Micro kit, Cat: 74004. RNA was processed at the University of Michigan Advanced Genomics Core to identify differential gene expression sets. DEG Pathway analysis was done using Advaita iPathway tools.

To test MHC-I expression cells were stained with anti-mouse H-2Kd-PE/Cy7 (Biolegend, Clone: SF1-1.1, RRID: AB_2562733, Cat: 116622) and anti-mouse H-2Dd-Alex fluorophore 647 (Biolegend, Clone: 34-2-12, RRID: AB_492913, Cat: 110612), anti-mouse CD21-PE (Biolegend, Clone 7E9, RRID: AB_940411 Cat: 123409), and goat anti-mouse C3d-FITC. Anti-C3d antibody was conjugated using Abcam FITC lightning link kit (Abcam, Cat: ab102884). Stained cells were analyzed by flow cytometry according to standard procedures in the laboratory^49^.

### RT-PCR

RNA was thawed on ice and concentration measured by nanodrop. 1.5μg RNA was brought up to 90μl with RNAse free water. cDNA was synthesized according to manufacturers’ protocol (Invitrogen, Superscript^TM^ VILO^TM^ cDNA synthesis kit, cat: 11754050) including an RT-negative control for each sample and a water only control in the RT+ and RT-reactions. cDNA was stored at -20°C. PCR PCR was performed using the following settings: (i) 50°C, 2 minutes; (ii) 95°C, 23 seconds; (iii) 60°C, 30 seconds. (ii) and (iii) were repeated for 40 cycles. Primers and probes were purchased from TaqMan and are shown in table labelled ST-primers.

### Immunofluorescence

Cells were plated at 1×10^6^, 5×10^5^, and 2.5×10^5^cells/100ml in separate wells in a U-bottom 96 well and incubated for one hour at 37°C. Following, the incubation, cells were treated with 200 μg/ml of C3d with an internalization domain, commercial C3d without an internalization domain, (R&D cat:2655-C3-050), or an equivalent volume of PBS supplemented with 10% glycerol for one hour at 37°C. Cells were harvested, washed in PBS, and centrifuged onto a microscope slide at 1500 rpm for five minutes using a Thermo-Scientific Shandon Cytospin 4. Cells were fixed with 4% paraformaldehyde for ten minutes at room temperature, washed three times with PBS, and permeabilized with 0.1% Triton X for 5 minutes at room temperature. Slides were then washed three times with PBS-Tween 20 (0.01%). Slides were incubated overnight at 4°C with goat IgG anti-mouse C3d antibody, (R&D Systems Cat: AF2655), diluted 1:125 in PBST containing 0.1% BSA. Slides were washed three times with PBST and incubated for one hour at 4°C with donkey anti-goat IgG-CF555 conjugated (Biotium-Fisher scientific, Cat: 50-196-4162), diluted 1:100 in PBST containing 0.1% BSA. Following the final three washes in PBST, nuclei were stained with Invitrogen’s ProLong TMGold anti-fade DAPI reagent (Cat: P36935), and slides were mounted. Slides were stored at -20°C until imaged. Imaging: DAPI and C3d stained images were obtained with a gain of 5.00, gamma set to 1.00 and exposure set to 750ms. All phase contrast images were taken with an exposure of 6ms, gain of 3, gamma set to 1.00. Photoshop: The intensity level of the C3d images were adjusted to the same degree using Adobe Photoshop 24.1.1. The levels of DAPI and phase contrast images were adjusted to similar brightness.

#### Data and Code Availability

Original/non-modified data will be stored in GEO and code for analyzing the data and creating figures are found will be deposited in Github upon manuscript acceptance.

### Statistics

Statistical analysis was performed using GraphPad Prism software (Version 10.2.3, Mac Version). Distributions of 2 unpaired groups were compared using two non-parametric two tailed Mann-Whitney tests. Paired analysis was by either multiple paired T tests or by the Wilcoxon matched-pairs signed rank test. Correction for multiplicity of measurements was done using False Discovery Rate.

Comparisons of more than 2 groups were by ANOVA. Categorical variables were compared using χ2 or Fisher’s exact test analysis. A P value of less than 0.05 was considered statistically significant. Data are presented as the mean ± SEM in all figures in which error bars are shown.

## Supporting information

Primers

## Funding

The research was supported by:

Grant from the State of Michigan Economic Development Corporation, CASE-283529 UM-MTRAC for Life Sciences (MC, JLP)

Department of Defense grant W81XWH-18-1-0721 (MC, JLP).

National Institutes of Health:

P01 CA093900 (ETK) supporting the Cancer Center shared resources (Single Cell and Spatial Analysis and Cancer Data Sciences Shared Resources),

P30 CA046592 (supports Cell and tissue imaging at the University of Michigan; PD Benjamin Allen) P30-AG024824 (ST)

The research was also supported by computational resources and services provided by Advanced Research Computing (ARC), a division of Information and Technology Services (ITS) at the University of Michigan, Ann Arbor.

The authors would like to acknowledge Dr. Bergsagel (Mayo Clinic, Arizona) who kindly provided the mouse models, read and edited the manuscript.

Declaration of Interests:

Drs. Platt and Cascalho are the inventors in two patents concerning C3d uses in Cancer. PCT NO:US-01/34733-converted and awarded in 2021, and U.S. Prov Pat Appl 63459844

## Author Contributions

JLP and MC conceived the research, analyzed data and wrote the manuscript. CZ performed most of the experiments and was helped by MGB, JH, JC and TPA. Analysis of single cell RNA data was by ST, AR and EK.

Analysis of Ig sequence data was by SS and MC. MP, EF, LN and LA provided useful feedback on the analysis of data and interpretation of findings. EF performed the pathology analysis of kidney sections.

Conceptualization: JLP and MC

Methodology: CZ, MGB, JH, JC and TPA

Investigation: SBB, DLA, MPW, WCB

Visualization: SFB, MJM, JLS, EH, ST, AR and EK

Funding acquisition: MC and JLP

Project administration: JLP and MC

Supervision: CZ, JLP and MC

Writing – original draft: JLP and MC

Writing – review & editing: JLP, MC, MP, EF, LN and LA

## List of Supplementary Materials

Figures S1 to S8, supplementary figures.

Data files ST-primers and ST1 to ST12 (csv files)

## Data and materials availability

All data, code, and materials used in the analysis will be made available to any researcher for purposes of reproducing or extending the analysis. There are no restrictions on materials with the exception of patent covered C3d recombinant peptide. Sequence data and code will be deposited in a public database. Those data include the scRNA data obtained from the plasma cells from mice treated with and without C3d, code to separate malignant from non-malignant plasma cells. All other data are available in the main text or the supplementary materials.”

