## Supplemental Figures for "Complement C3d enables protective immunity capable of distinguishing spontaneously transformed from non-transformed cells"

**Complement-dependent cell-mediated immunity eliminates multiple myeloma malignant clones by enhancing antigen presentation**

**Authors:** Jeffrey L Platt[1,2*], Chong Zhao[1,2], Jeffrey Chicca[1], Matthew Pianko[3], Joshua Han[1,2], Stephanie The[4,5], Arvind Rao[4,5], Evan Keller[6], Mayara Garcia de Mattos Barbosa[1], Lwar Naing[1,8], Tracy Pasieka-Axenov[1], Lev Axenov[1], Simon Schäfer[9], Evan Farkash[7], and Marilia Cascalho[1,2,*].

**Extended data figures:**

**Figure S1 A**. C3d peptide sequence and purification. The C3d peptide sequence encompasses amino acids 1024 to 1320 of the C3d region of murine C3. The sequence includes a N-terminal protein transduction domain (RKKRRQRRR) and Cys (C) to Ser(S) substitution at position 1028 of C3 to avoid the presence of a free sulfhydryl in the recombinant peptide. The C3d sequence was cloned into a baculovirus expression vector and used to transduce Sf9 insect cells. Large-scale production for purification was done in High-five infected at an MOI of 2. Purification was done using NTA Agarose from Qiagen and eluted with imidazole in PBS. The samples were analyzed on a 4-20% gradient SDS-PAGE. The monomer was detected in the soluble (shown) and in the insoluble fractions (not shown). Aliquots of 1mg/ml were prepared and mice were injected with 20 microliters, containing 20 micrograms of peptide in PBS intra tibia. **B**. Detection of M protein. Vk-MYC* mice were injected with C3d or PBS. Two weeks later, mice treated with C3d were either injected with anti-CD8 antibodies or with isotype control three times a week starting two weeks after C3d injection and continued for 6 weeks until experiment end. M protein was detected by SPEP before and at the end of the experiment. M protein seen as a single narrow band in the gamma region of the gel decreases in mice treated with C3d. Instead, a broad band of polyclonal Ig appears. In untreated mice and in mice treated with C3d followed by anti-CD8 antibodies the narrow band increases in intensity relative to the albumin band and remains narrow. **C.** Graph shows the relative change in M protein intensity, following each treatment and relative to albumin. M protein density was measured using image J.

**Figure S2: C3d decreases tumor load without changing other cell populations in the BM**. Vk*myc mice older (late disease) and younger (early disease) than one year of age, with M-protein, were injected with 20 microgram C3d peptide intra-tibia two months before analysis. Vk*myc were immunized twice, 2 months apart before treatment with C3d, M protein was detected by SPEP. **A and B.** Tsne plots of mass cytometry (CYTOF) indicating plasma cell clusters in purple. **C.** The graph depicts frequencies of plasma cells in the treated or control mice divided according to age at beginning of treatment assessed by mass cytometry. **D.** Analysis of the frequency of non-plasma cell bone marrow cell populations by mass cytometry. Frequencies of the total number of cells are indicated. Data analysis was by Mann-Whitney test. Comparisons that yielded $P<0.05$ are noted by **, P=0.0022; *, P=0.0286. Statistics analysis was by Prism software (v9 for mac).

**Figure S3: UMAP plots of sc RNA gene expression depicting the different cell populations**. A. UMAP plots of every sample considered in the analysis. B. Gene expression of markers used to differentiate between cell populations. Y-axis depicts the names of the cell types; the X-axis depicts the gene identity. Expression is noted according to the number of cells that express the marker above background (diameter of the circle increases with the fraction of cells expressing the marker). The average intensity is represented by a color gradient varying from white (no expression) to purple (mid intensity expression) and to blue (maximum expression).

**Figure S4**. **IgVH gene frequencies in B cells in the bone marrow of Vk-MYC* mice treated (red) or not (black) with C3d.** Differences in frequency according to treatment did not reach significancy by multiple Mann-Whitney tests.

**Figure S5: C3d treatment induces expression of genes encoding ribosomal proteins and LncRNAs *in vivo* and in B cells in culture. B cell lymphoma cells (**18.81) (1) ($10^5$/ml) were incubated with 20μg C3d, with PBS for 24h, 48h and 72h and RNA obtained and sequenced. For immunofluorescence, $10^6$ cells were incubated with 200 μg of C3d with an internalization domain (ID) or with C3d lacking ID. **A.** Immunofluorescence analysis of 18.81 B cells in cytospin slides stained with goat anti-mouse C3d followed by Texas-red labelled rabbit anti-goat antibodies. Bar indicate 50 microns. C3d with ID is incorporated by lymphoma B cells after one hour incubation while C3d without ID accumulates on the cell surface. **B.** C3d induced differential expression of genes in plasma cells of bone marrow of treated mice relative to untreated mice. Figure shows DEGs that that are also induced by C3d on 18.81 cells relative to untreated cells, after 48 hours incubation. Values are expressed as $Log_2$ Fold Change. Negative values mean decreased expression. Positive values mean increased expression. Changes in expression considered had an adjusted P <0.001 (ST6 and ST7). **C.** Volcano plot analysis of differential gene expression by 18.81 B cells incubated with C3d for 48 hours. In red are depicted all the differentially expressed lncRNAs. C3d changed the expression of 1093 genes with P<0.001 relative to untreated cells (ST7). **D.** Top 20 lncRNAs increased in 18.81 cells incubated with C3d for 48 hours in comparison with untreated cells.

**Figure S6: E2f1 Pathway is induced by C3d.** Lymphoma 18.81 cells (1) ($10^5$/ml) were incubated with 20μg C3d or with PBS for 24h, 48h and 72h. Following incubation cells were washed, RNA produced, sequenced and processed at the University of Michigan Advanced Genomics Core. **A.** Schematics of E2f1 pathway analyzed by String (https://string-db.org). Shown are genes differentially expressed by C3d relative to controls following C3d treatment of 18.81 B cells for 48 hours culture (P>0.05). Network nodes represent proteins. Colored nodes refer to first shell of interactor**s.** Edges represent protein interactions, known from curated data bases or experimental data; predicted by gene neighborhood, co-occurrence, homologies and/or co-expression. Software available at string-db.org/cgi run at the highest confidence (0.900) to show a maximum of 5 interactions and analysis with DBSCAN clustering to show a maximum of three clusters (String, version 12.0, String consortium 2023, used under a Creative Commons BY 4.0 license). **B.** Heat map of E2f1 pathway differentially expressed by 18.81 cells incubated with C3d for 48 hours relative to controls. **(C)** RT-PCR analysis of E2f1 pathway gene expression in RNA obtained from lymphoma cell cultured with or without C3d for 24 or 48 hours.

**Figure S7: C3d treatment increases MHC class-I expression by 18.81 cells. A-B.** MHC class I gene expression at 24h, 48h and 72h after incubation with C3d or in control cultures. H-2Kd expression was analyzed by FACS. Plots depict live cells gated on lymphocytes. Shown in **(A)** is H-2Kd MFI at 48h of culture in non-dividing cells. Shown in **(B)** is the graphic representation of the MFIs of three replicate cultures with C3d or untreated after, 24 or after 48 and 72 hours. Statistical analysis was by ANOVA followed by Kruskal-Wallis test (Prism for Mac v10.2.3).

**Figure S8: Differential gene expression in the bone marrow of mice treated with C3d relative to untreated mice**. Volcano Plot on the left depicts the top DEG expressed by bone marrow cells annotated according to the cell of origin. Volcano plot on right depicts DEG expressed by Plasma Cells (PCs) in the bone marrow cells of mice treated with C3d or left untreated. Positive fold changes correspond to genes that increase in expression by C3d while negative fold changes correspond to genes that decrease in expression following C3d treatment. Changes in expression that had an adjusted P <0.05 were included. Data plotted can be found in ST6.

**Supplementary Data Tables:**
ST-primers. Taq Man primers and probes used to detect E2f1 pathway gene expression.

ST 1. Single cell RNA gene expression per cluster comparing each cluster to all the others to differentiate clusters, indicating the $Log_2$ of the fold change and adjusted P in all the mice.

ST 2. V(D)J clonotype frequencies. A clonotype was defined as >80% CDR 3 AA sequence overlap with same VH and JH heavy chain genes.

ST 3. Expression of malignancy associated genes by each cell to define malignant and non-malignant clones.

ST 4. Principal Component Analysis based on expression of malignancy associated genes.

ST 5. Malignancy and non-malignancy scores per cell.

ST 6. Single cell RNA differential gene expression by every cell cluster comparing cells obtained from the bone marrow of C3d treated mice with cells obtained from untreated mice. Genes that are differentially expressed with an Adj P<0.001 are highlighted in yellow. A volcano plot shown in figure S8 illustrates differentially expressed genes in every cluster noting the 20 top C3d-induced genes. A separate plot is shown for genes differentially expressed by Plasma Cells (Figure S8).

ST 7. Differentially expressed genes after 48 hours incubation of 18.18 pre-B cells with C3d or glycerol. Positive fold change values mean increased expression in C3d treated cells; negative values mean decreased expression in C3d treated cells. Genes that are differentially expressed with an Adj P<0.001 are highlighted in yellow.

ST 8. Differential LncRNA gene expression after 48 hours incubation of 18.18 pre-B cells with C3d or glycerol.

ST 9-10. Single Cell RNA differential gene expression by T cells from C3d treated mice in comparison to T cells obtained from the bone marrow of mice not treated with C3d. $Log_2$ transformed fold change and adjusted P are indicated. Positive values mean increased expression in C3d treated mice; negative values mean decreased expression in C3d treated mice. ST10 data is formatted for Volcano plot display.

ST 11-12. Single Cell RNA differential gene expression by granulocytes obtained from the bone marrow of C3d treated mice in comparison to granulocytes obtained from the bone marrow of mice not treated with C3d. $Log_2$ transformed fold change and adjusted P are indicated. Positive values mean increased expression in C3d treated mice; negative values mean decreased expression in C3d treated mice. ST12 data is formatted for Volcano plot display.

A

**Purified recombinant C3d attached to a protein transduction domain (RKKRRQRRR)**

```
        10         20         30         40         50
TPAGSGEQHM IGMTPTVIAV HYLDQTEQWG KFGIEKRQEA LELIKKGYTQ
        60         70         80         90        100
QLAFKQPSSA YAAFNNRPPS TWLTAYVVKV FSLAANLIAI DSHVLCGAVK
       110        120        130        140        150
WLILEKQKPD GVFQEDGPVI HQEMIGGFRN AKEADVSLTA FVLIALQEAR
       160        170        180        190        200
DICEGQVNSL PGSINKAGEY IEASYMNLQR PYTVAIAGYA LALMNKLEEP
       210        220        230        240        250
YLGKFLNTAK DRNRWEEPDQ QLYNVEATSY ALLALLLLKD FDSVPPVVRW
       260        270        280        290
LNEQRYYGGG YGSTQATFMV FQALAQYQTD VPDHKDLNMD VSFHLPSRS
```

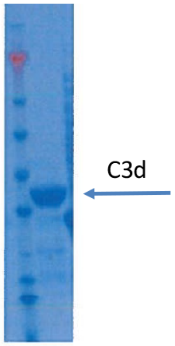


MW = 35.1 kDa
1mg/ml in PBS

B

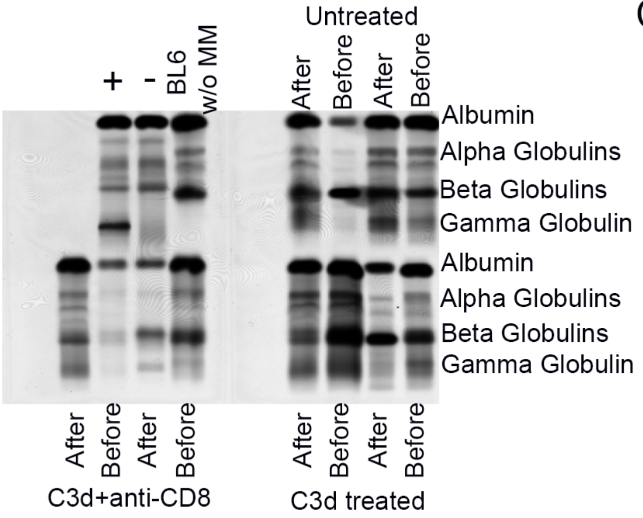


C

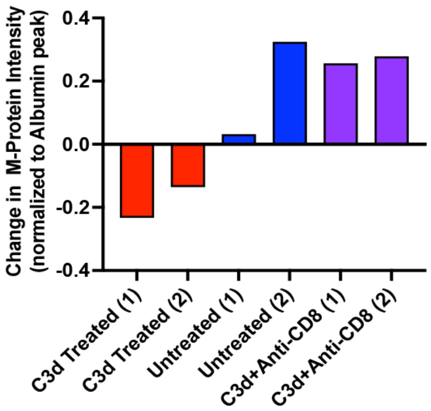


Figure S1

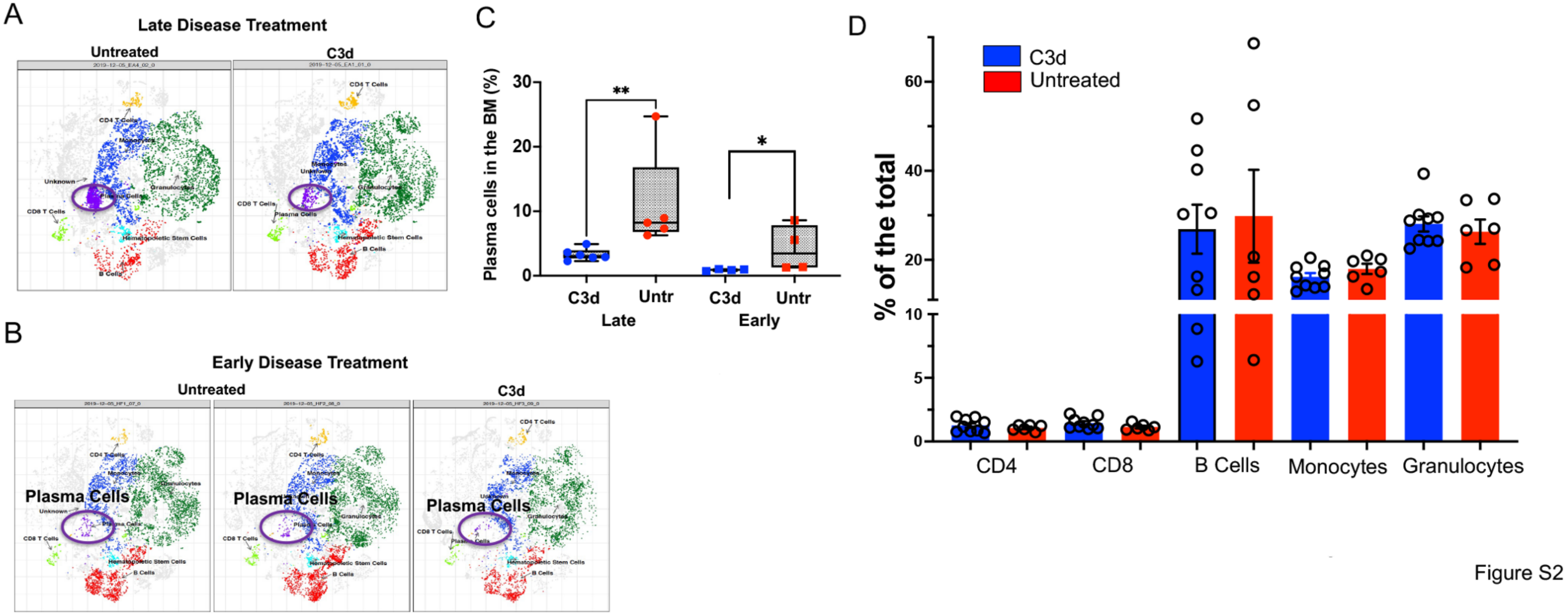
A
Late Disease Treatment
Untreated
C3d
B
Early Disease Treatment
Untreated
C3d
Plasma Cells
Plasma Cells
Plasma Cells
C
Plasma cells in the BM (%)
**
*
C3d
Untr
C3d
Untr
Late
Early
D
C3d
Untreated
% of the total
CD4
CD8
B Cells
Monocytes
Granulocytes
Figure S2

Figure S2

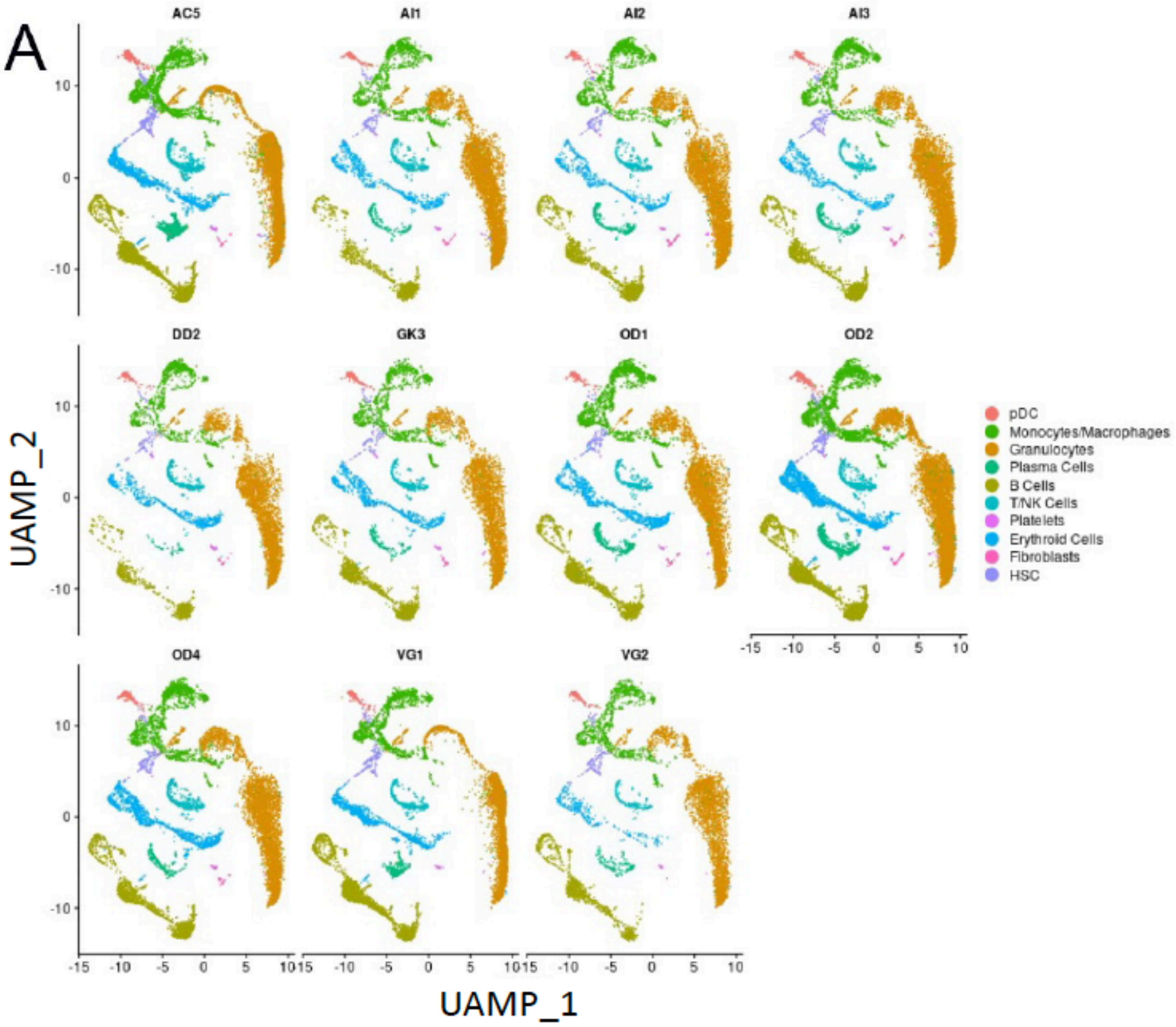

A
AC5
AI1
AI2
AI3
DD2
GK3
OD1
OD2
OD4
VG1
VG2
UAMP_2
UAMP_1
pDC
Monocytes/Macrophages
Granulocytes
Plasma Cells
B Cells
T/NK Cells
Platelets
Erythroid Cells
Fibroblasts
HSC


Figure S3A

# B

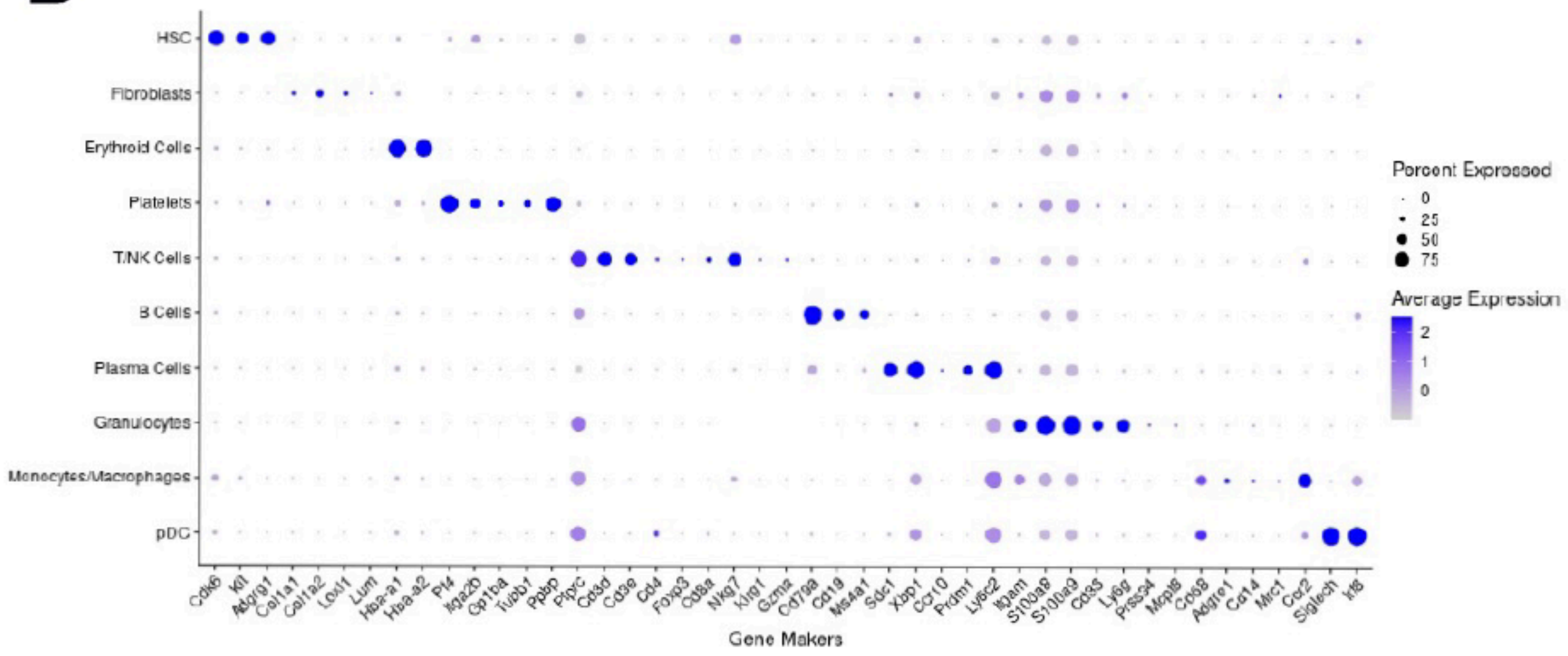
HSC
Fibroblasts
Erythroid Cells
Platelets
T/NK Cells
B Cells
Plasma Cells
Granulocytes
Monocytes/Macrophages
pDC
Cdk6
Kit
Adgrg1
Col1a1
Col1a2
Loxl1
Lum
Hba-a1
Hba-a2
Pf4
Itga2b
Gp1ba
Tubb1
Ppbp
Ptprc
Cd3d
Cd3e
Cd4
Foxp3
Cd8a
Nkg7
Klrg1
Gzma
Cd79a
Cd19
Ms4a1
Sdc1
Xbp1
Ccr10
Prdm1
Ly6c2
Itgam
S100a8
S100a9
Cd33
Ly6g
Prss34
Mcpt8
Cd68
Adgre1
Cd14
Mrc1
Ccr2
Siglech
Irf8
Gene Makers
Percent Expressed
0
25
50
75
Average Expression
2
1
0

Figure S3B

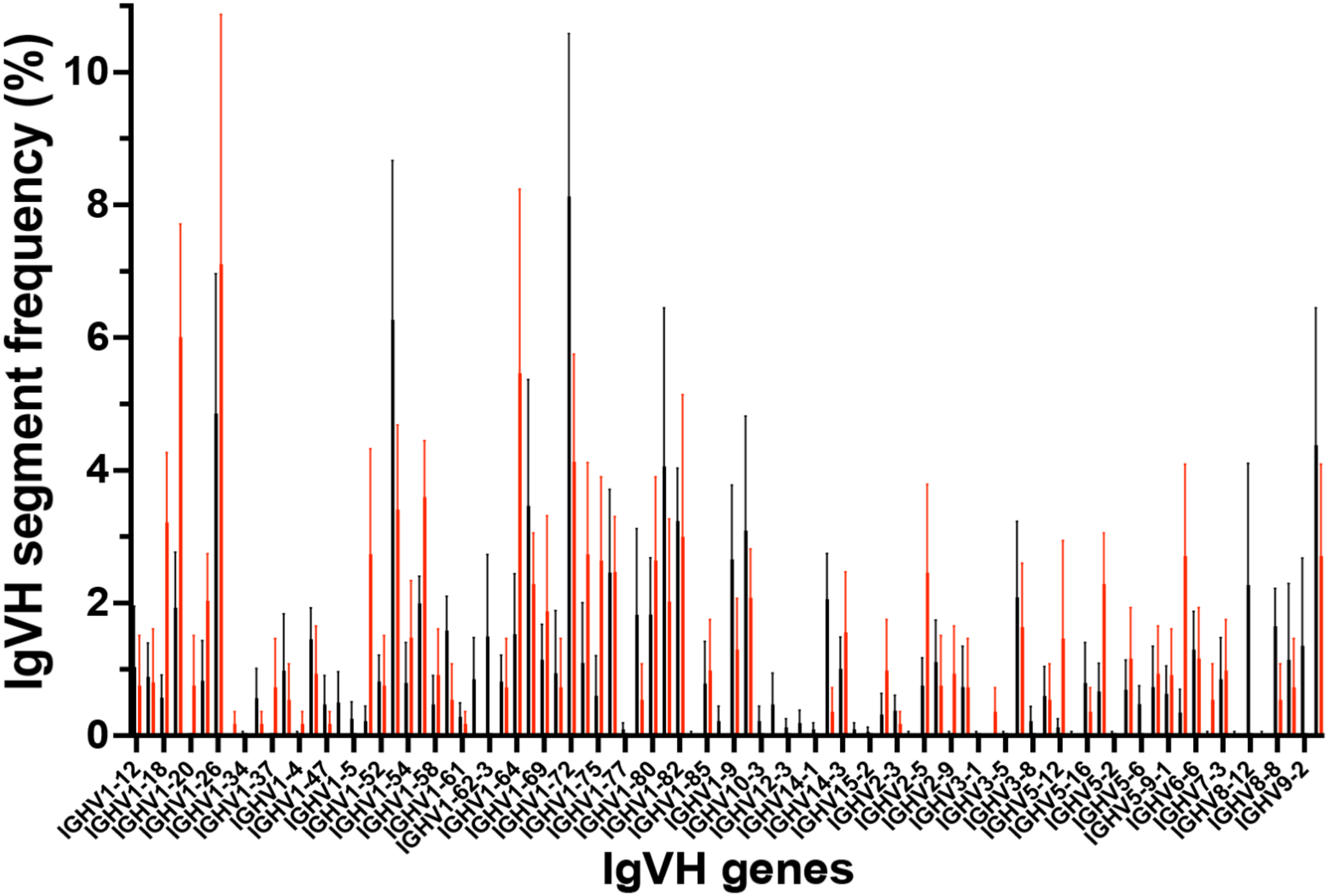
IgVH segment frequency (%)
0
2
4
6
8
10
IGHV1-12
IGHV1-18
IGHV1-20
IGHV1-26
IGHV1-34
IGHV1-37
IGHV1-4
IGHV1-47
IGHV1-5
IGHV1-52
IGHV1-54
IGHV1-58
IGHV1-61
IGHV1-62-3
IGHV1-64
IGHV1-69
IGHV1-72
IGHV1-75
IGHV1-77
IGHV1-80
IGHV1-82
IGHV1-85
IGHV1-9
IGHV10-3
IGHV12-3
IGHV14-1
IGHV14-3
IGHV15-2
IGHV2-3
IGHV2-5
IGHV2-9
IGHV3-1
IGHV3-5
IGHV3-8
IGHV5-12
IGHV5-16
IGHV5-2
IGHV5-6
IGHV5-9-1
IGHV6-6
IGHV7-3
IGHV8-12
IGHV8-8
IGHV9-2
IgVH genes

Figure S4

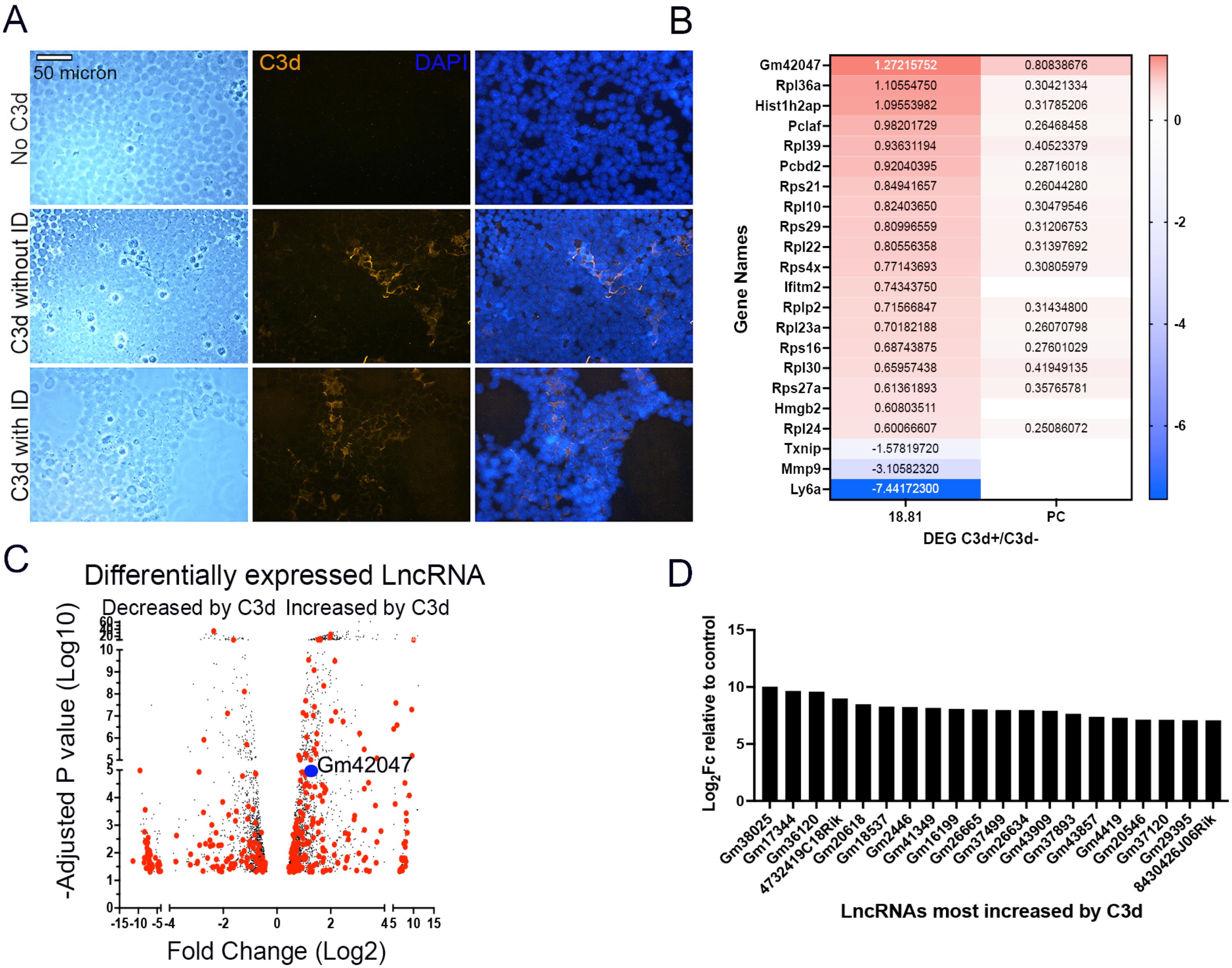
A
50 micron
C3d
DAPI
No C3d
C3d without ID
C3d with ID
B
Gene Names
Gm42047 1.27215752 0.80838676
Rpl36a 1.10554750 0.30421334
Hist1h2ap 1.09553982 0.31785206
Pclaf 0.98201729 0.26468458
Rpl39 0.93631194 0.40523379
Pcbd2 0.92040395 0.28716018
Rps21 0.84941657 0.26044280
Rpl10 0.82403650 0.30479546
Rps29 0.80996559 0.31206753
Rpl22 0.80556358 0.31397692
Rps4x 0.77143693 0.30805979
Ifitm2 0.74343750
Rplp2 0.71566847 0.31434800
Rpl23a 0.70182188 0.26070798
Rps16 0.68743875 0.27601029
Rpl30 0.65957438 0.41949135
Rps27a 0.61361893 0.35765781
Hmgb2 0.60803511
Rpl24 0.60066607 0.25086072
Txnip -1.57819720
Mmp9 -3.10582320
Ly6a -7.44172300
18.81
PC
DEG C3d+/C3d-
0
-2
-4
-6
C
Differentially expressed LncRNA
Decreased by C3d
Increased by C3d
-Adjusted P value (Log10)
Gm42047
Fold Change (Log2)
D
Log₂Fc relative to control
Gm38025
Gm17344
Gm36120
4732419C18Rik
Gm20618
Gm18537
Gm2446
Gm41349
Gm16199
Gm26665
Gm37499
Gm26634
Gm43909
Gm37893
Gm43857
Gm4419
Gm20546
Gm37120
Gm29395
8430426J06Rik
LncRNAs most increased by C3d

Figure S5

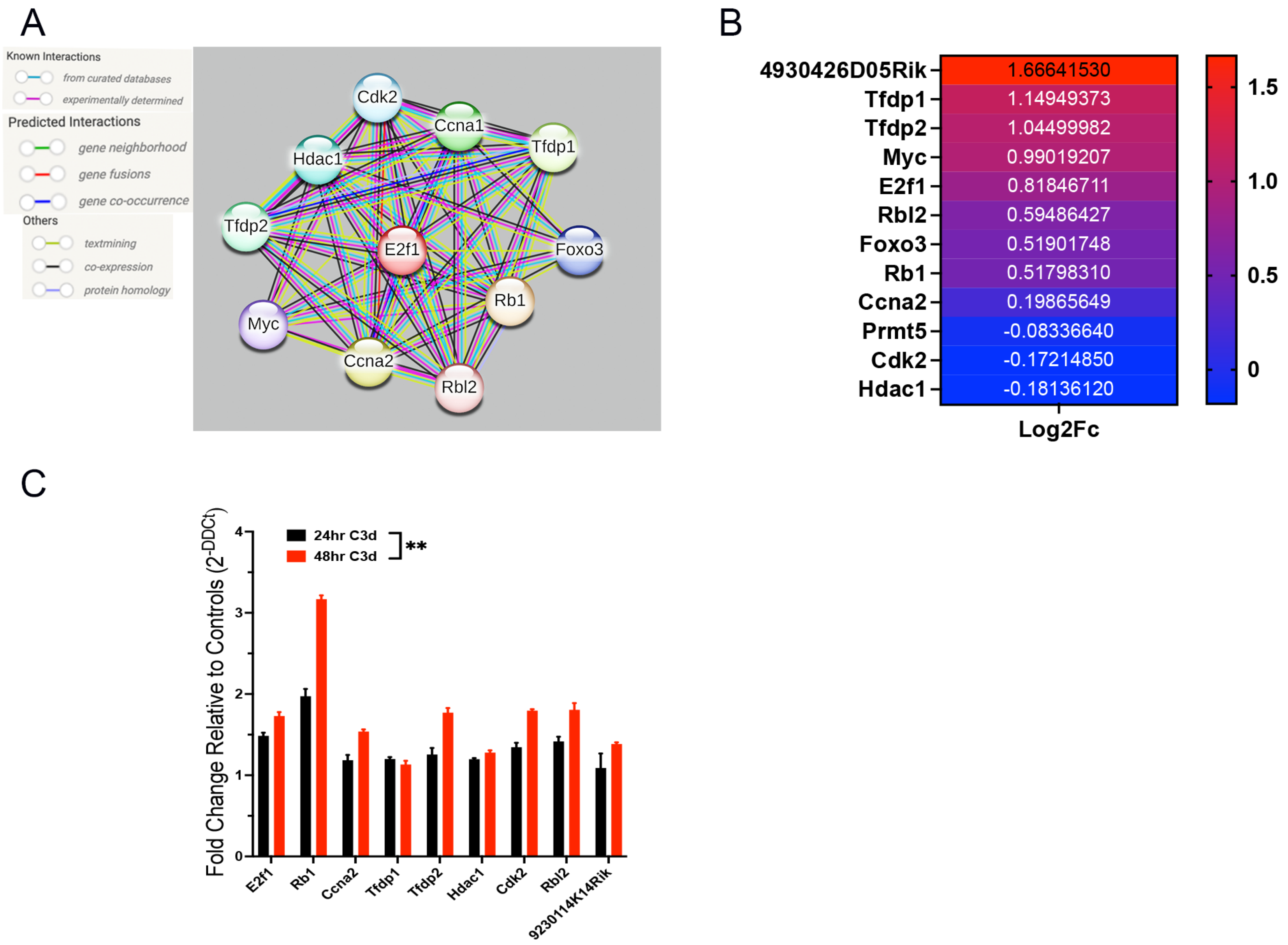
A
Known Interactions
from curated databases
experimentally determined
Predicted Interactions
gene neighborhood
gene fusions
gene co-occurrence
Others
textmining
co-expression
protein homology
Cdk2
Ccna1
Tfdp1
Hdac1
Tfdp2
E2f1
Foxo3
Rb1
Myc
Ccna2
Rbl2
B
4930426D05Rik 1.66641530
Tfdp1 1.14949373
Tfdp2 1.04499982
Myc 0.99019207
E2f1 0.81846711
Rbl2 0.59486427
Foxo3 0.51901748
Rb1 0.51798310
Ccna2 0.19865649
Prmt5 -0.08336640
Cdk2 -0.17214850
Hdac1 -0.18136120
Log2Fc
1.5
1.0
0.5
0
C
Fold Change Relative to Controls (2-DDCt)
24hr C3d
48hr C3d
**
0
1
2
3
4
E2f1
Rb1
Ccna2
Tfdp1
Tfdp2
Hdac1
Cdk2
Rbl2
9230114K14Rik

Figure S6

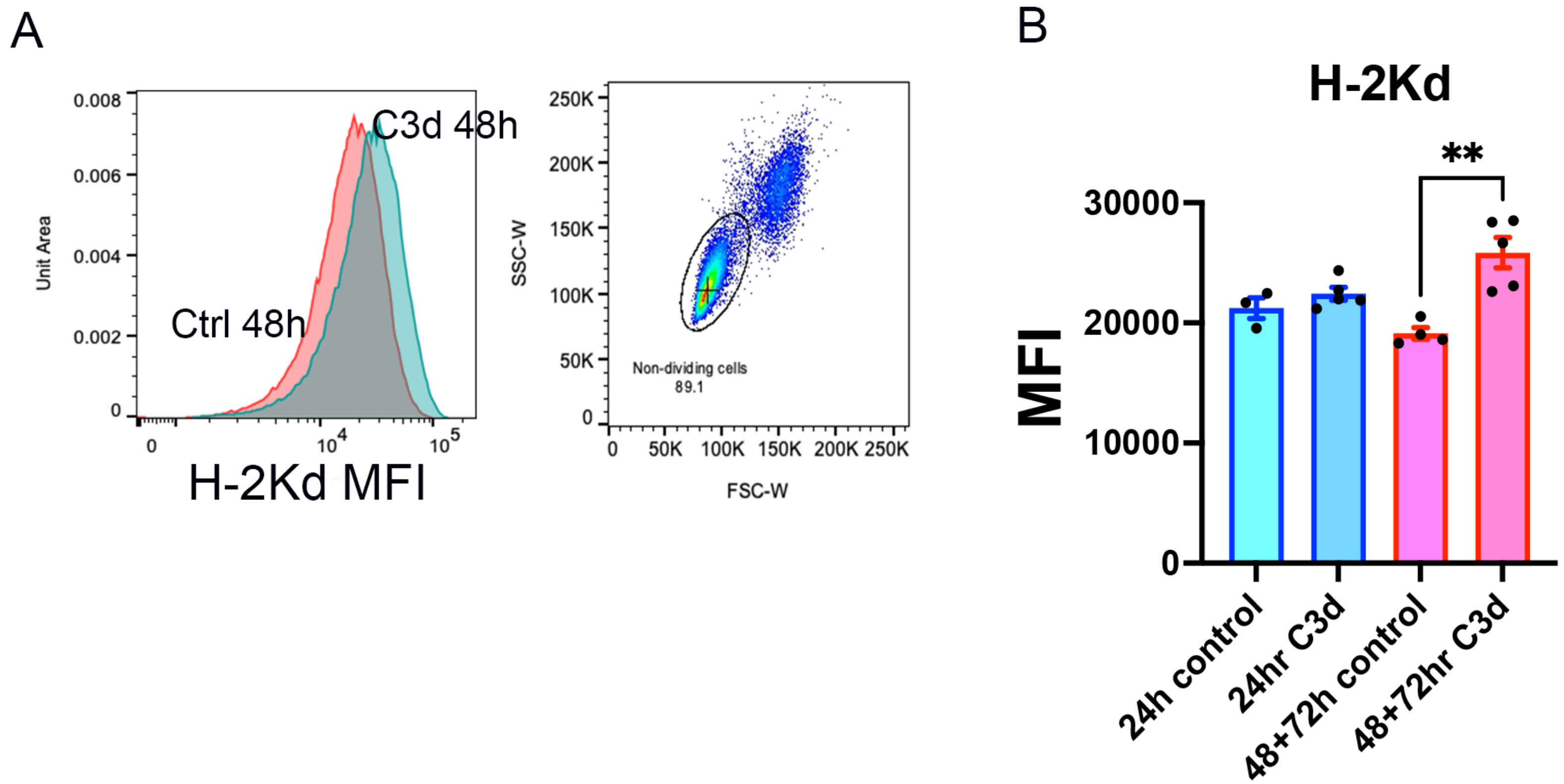
A
0.008
0.006
0.004
0.002
0
Unit Area
C3d 48h
Ctrl 48h
0
10^4
10^5
H-2Kd MFI
250K
200K
150K
100K
50K
0
SSC-W
Non-dividing cells
89.1
0
50K
100K
150K
200K
250K
FSC-W
B
H-2Kd
**
30000
20000
10000
0
MFI
24h control
24hr C3d
48+72h control
48+72hr C3d

Figure S7

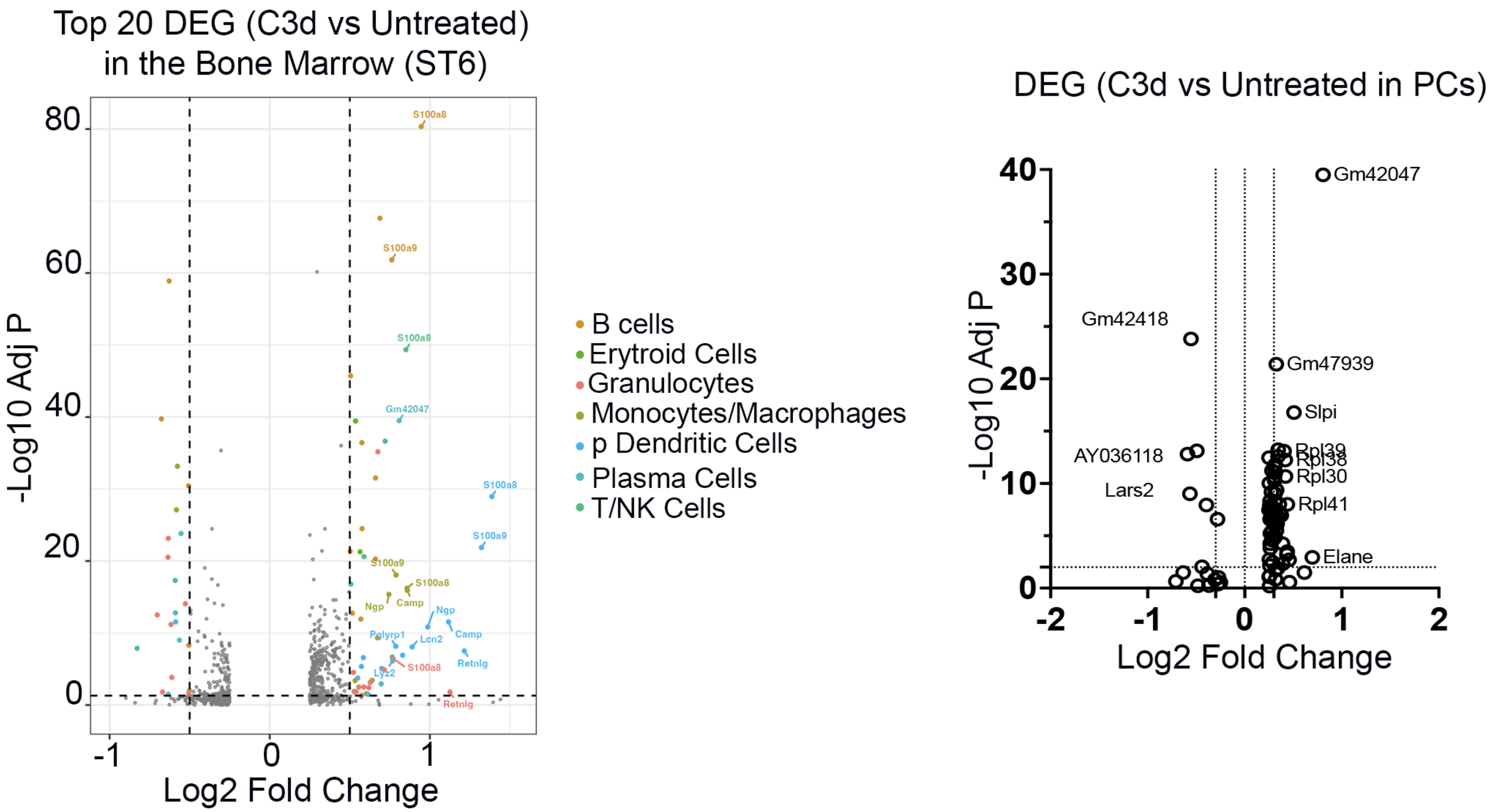
Top 20 DEG (C3d vs Untreated)
in the Bone Marrow (ST6)
-Log10 Adj P
Log2 Fold Change
S100a8
S100a9
Gm42047
Ngp
Camp
Pglyrp1
Lcn2
Retnlg
Lyz2
B cells
Erytroid Cells
Granulocytes
Monocytes/Macrophages
p Dendritic Cells
Plasma Cells
T/NK Cells
DEG (C3d vs Untreated in PCs)
-Log10 Adj P
Log2 Fold Change
Gm42047
Gm42418
Gm47939
Slpi
AY036118
Rpl39
Rpl38
Rpl30
Lars2
Rpl41
Elane

Figure S8
